# Outer Pore Collapse as a Potential Mechanism of Partial Loss of Pain in Nav1.7 M899I

**DOI:** 10.64898/2026.08.18.745213

**Authors:** Vishal Sudha Bhagavath Eswaran, Estefania Torres-Ortiz, Petra Hautvast, Amdiya Botchoi, Silvia Detro-Dassen, Anika Neureiter, Yi Liu, Ralf Hausmann, Angelika Lampert

## Abstract

Complete loss of function of the voltage-gated sodium channel subtype Na_v_1.7, encoded by SCN9A, results in congenital insensitivity to pain. Here, we investigate a previously identified variant, M899I, in which methionine at position 899 is substituted by isoleucine. This variant was originally described in a Chinese patient with loss of pain. We confirmed membrane expression of the mutant channel in HEK cells using extracellular HA-tagging; however, no sodium currents were detectable from the variant in patch-clamp recordings.

The M899I substitution is located within a tightly packed hydrophobic region of the pore module. Introducing the corresponding variant into Na_v_1.2 and Na_v_1.5 similarly abolished channel function, underscoring the high conservation and functional importance of this residue. To further investigate the underlying mechanism, we combined *in-silico* coarse-grained molecular dynamics simulations with *in-vitro* electrophysiological analysis. Our simulations predicted that the M899I substitution induces collapse of the outer pore, substantially reducing both pore radius and volume. Substitution with other hydrophobic residues was likewise predicted to alter pore geometry and, consequently, ion permeation to varying degrees. Whole-cell voltage-clamp recordings validated these predictions, with observed current densities closely correlating with the extent of pore collapse predicted *in silico*.

Together, our findings establish pore collapse as a mechanism underlying disease-relevant loss-of-function variants in Na_v_1.7 and suggest that this principle may extend to other sodium channel subtypes. Moreover, our results demonstrate that *in-silico* molecular dynamics approaches can reliably predict structural and functional consequences of channel variants, as confirmed by *in-vitro* electrophysiological data.

**Statement of Significance:** Na_v_1.7 is a key determinant of pain perception, and genetic variants in this channel are known to cause a spectrum of pain disorders. Understanding the structural and functional consequences of these variants is essential for elucidating the mechanisms that govern channel function.

Using complementary *in silico* and *in vitro* approaches, we demonstrate that the loss-of-function Na_v_1.7 variant p.M899I, associated with congenital pain insensitivity, is expressed at the plasma membrane but induces alterations in pore geometry. Our findings underscore the critical role and high sensitivity of the pore module in ion conduction and suggest a broader pathogenic mechanism that may be shared among loss-of-function Na_v_ channel variants affecting the outer pore region.

## Introduction

Voltage-gated sodium channels (Na_v_s) are transmembrane proteins that play a critical role in the generation and propagation of action potentials. Of the nine subtypes (Na_v_1.1-Na_v_1.9) expressed in humans, Na_v_1.7 has been shown to play a significant role in signalling of potentially painful stimuli (1). The α-subunit of Na_v_1.7, and all eukaryotic Na_v_s in general, are folded into four homologous domains from a single polypeptide (2). Each domain is made of six transmembrane helices (S1-S6), two membrane re-entrant pore helices (P1 and P2) and a helical segment connecting S4 and S5 (S4-S5 linker) (Figure 1A). The domains are connected by long intracellular linkers (D1-D2, D2-D3 and D3-D4). The S1-S4 helices form the voltage sensing domain (VSD), while the S5-S6 and P1-P2 helices form the pore module (PM). The PM is important for the selectivity to and conduction of sodium ions through the pore, with the ‘DEKA’ motif – a set of four amino acids - acting as the selectivity filter (2). The α-subunit can also interact with secondary proteins like the β subunits (ß1-ß4) that either interact via covalent disulfide bonds (ß2 and ß4) or via non-covalent interactions (ß1 and ß3) (2). Evidence from subtypes such as Na_v_1.5, Na_v_1.2, Na_v_1.8 and Na_v_1.7 have added to the notion that Na_v_s are likely to dimerize - whereby two α-subunits interact with each other - and also gate in a coupled manner (3–6).

**Figure 1:**
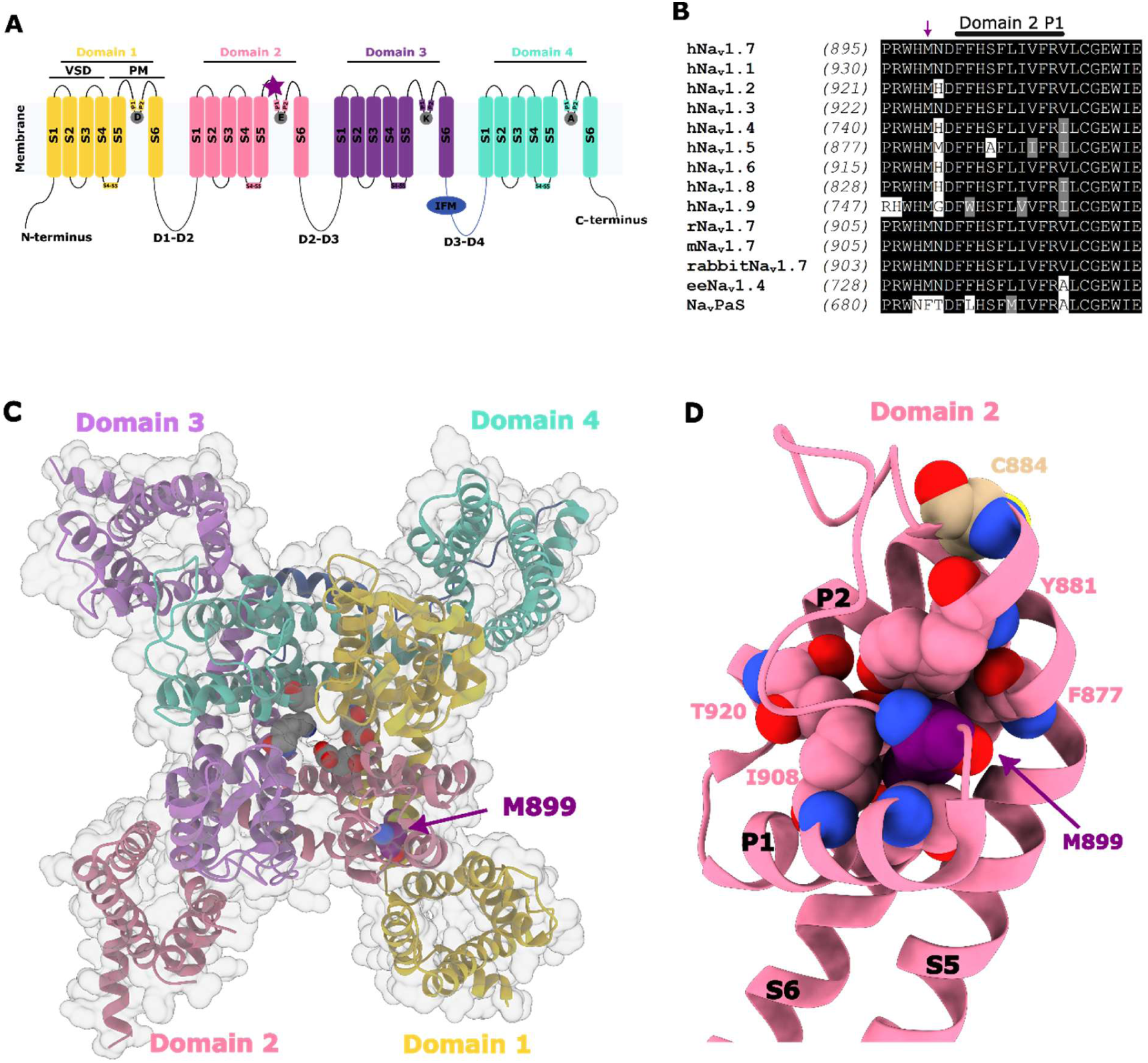
Structural location and sequence conservation of the M899 residue. **A)** 2-D schematic of a human voltage-gated sodium channel (Na_v_s), representing four domains (domain 1-4), each comprising six transmembrane helices (S1-S6) and two re-entrant pore helices (P1-P2). The IFM motif important for fast inactivation is shown in blue. The residue of interest M899 is located in the PM of domain 2 (purple star). **B)** M899 (purple arrow) is fully conserved across hNa_v_s and Na_v_s of various species, with the only exception being Na_v_Pas that contains a phenylalanine (F) instead of a methionine (M). Multiple sequence alignment of the nine human Na_v_ subtypes (hNa_v_1.1-1.9), Na_v_1.7 of other mammalian species (mouse mNa_v_1.7, rat rNa_v_1.7 and rabbit rabbitNa_v_1.7), Na_v_1.4 of the electric eel (eeNa_v_1.4) and the Na_v_ of the american cockroach (Na_v_PaS). Amino acids are represented by their single letter representation. Grey boxes show partially conserved residues and non-shaded boxes show non-conserved residues. The italicized number in brackets represent the starting number of each sequence region shown. **C)** A 3-D extracellular view of the hNa_v_1.7 cryo-EM structure in a fast-inactivated state (PDB ID: 6J8G) (36). The four domains are shown with a cartoon representation and color coded. The residue of interest M899 (purple sphere) is located in the PM of domain 2. M899 is close to the DEKA selectivity filter (grey spheres) located in the outer pore region. **D)** A close-up of the PM of domain 2 in cartoon representation. M899 (purple sphere) is tightly packed within a hydrophobic pocket and surrounded by various hydrophobic residues (pink spheres). M899 is close to the C894 residue (sand colored spheres) involved in binding the ß2 subunit.

Genetic variants in SCN9A, the gene encoding Na_v_1.7, have been linked to aberrations in pain sensation (2). The effect of these variants on Na_v_1.7 can broadly be categorized as gain of function (GoF) or loss of function (LoF). Whereas gain-of-function (GoF) variants are commonly associated with severe pain disorders, including inherited erythromelalgia and paroxysmal extreme pain disorder, loss-of-function (LoF) variants have been implicated in the loss of pain sensation characteristic of congenital insensitivity to pain (CIP) (7).

CIP-causing variants can either cause truncations that lead to non-functional and partially-folded Na_v_1.7 (8, 9), or alter a single or set of amino acid residues leading to disrupted membrane expression or gating of Na_v_1.7 (10, 11). In heterozygous individuals, loss-of-function (LoF) variants can dimerize with a functional wild-type (WT) channel and reduce channel activity by more than 80% when co-expressed at a 1:1 ratio. These so-called dominant-negative (DN) effects arise when the mutant channel impairs the function of its WT counterpart and have been reported for LoF Na_v_1.7 variants (5).

Understanding the effect of missense variants on the individual α-subunit, however, is a more complex problem, as the channel structure and gating must be taken into account. Since the publishing of the first cryo-EM structure of hNa_v_1.7 (12), advances in this field have led to an increasing number of structures being released that has helped us understand key concepts of Na_v_ channel structure and gating (G. Huang et al., 2022; Jiang et al., 2021; Jiang et al., 2020; Z. Li et al., 2021). The increasing availability of structural data has advanced molecular dynamics simulations, allowing detailed characterization of Na_v_ channel gating and the effects of mutations in physiologically relevant environments comprising lipids, water, and ions. Nevertheless, the consequences of patient-derived variants cannot be fully understood from structural analyses in isolation and require complementary experimental validation. Here, we investigate the variant SCN9A c.2697G>A (hNa_v_1.7 p.M899I) that was identified in a patient with a reduction in pain sensation (17). By applying both *in-silico* and *in-*vitro methodologies we predict the impact of the variant on pore geometry and function and its subsequent effect on channel conduction, highlighting the connection between pore module folding and conduction.

## Materials and Methods

### Cell culture and transient transfection

HEK293t cells were cultured in Dulbecco’s modified eagle medium F-12 (DMEM-F12; Thermo Fischer Scientific, Waltham, MA, USA) supplemented with 10% fetal bovine serum (FBS good; PAN-Biotech, Aidenbach, Germany) and incubated at 37°C with 5% CO_2_. The hNa_v_1.7 WT cDNA was contained in a pCMV6Neo vector (Origene, Rockville, MD, USA). The cDNA of the various substitutions of the M899 residue namely M899A, M899C, M899F, M899I, M899L and M899V generated by utilizing site-directed mutagenesis on the hNa_v_1.7 WT and contained in a pCMV6Neo vector (Origene, Rockville, MD, USA). The cDNA of the hNa_v_1.2 WT was contained in a pCMV6-XL5 vector (Gift from Frank Bosmans, Vrije Universiteit Brussel) with M925T introduced via site-directed mutagenesis of the hNa_v_1.2 WT cDNA and contained in the same vector. hNa_v_1.5 WT instead was contained in a pTracer™-SV40 vector (Thermo Fischer Scientific, Waltham, MA, USA). hNa_v_1.5 M881I was generated via site-directed mutagenesis of hNa_v_1.5 cDNA and contained in the same vector. The human ß2 subunit cDNA was contained in a pGEM vector (Thomas Zimmer, Uni Jena) (18).

Transient transfection of cultured HEK293t cells with the hNa_v_1.7 WT, hNa_v_1.2 WT, hNa_v_1.5 WT or any of the mutations of these three subtypes were done using JetPEI® (Polyplus-transfection S.A., Illkirch, France) according to the manufacturer’s protocol. Briefly, HEK293t cells were cultured in 35mm dishes 1 day before transfection. 1.25µg of the plasmid of interest along with 0.25µg of pMax-GFP (GFP) as a reporter protein was diluted in 50µL of 150mM NaCl solution. 3µL of JetPEI® reagent was buffered in 50µL of 150mM NaCl solution and this mixture was then added to the DNA mixture and gently mixed. The resulting mixture was allowed to incubate at room temperature for 15-20 minutes before being added to the cells. For experiments involving the human ß2 subunit, the 1.5µg of DNA consisted of 1.25µg hNa_v_1.7 M899I, 0.125µg GFP and 0.125µg of human ß2. The rest of the protocol remained unaltered. The transfected cells were incubated overnight at 37°C and 5% CO_2_. For experiments to test if temperature can alter hNa_v_1.7 M899I expression, cells were incubated overnight at 30°C and 5% CO_2_. For experiments involving co-expression of hNav1.7 M899I and hNa_v_1.7 WT, both plasmids were transfected to the same cells in five different ratios – 1:0, 1:4, 1:1, 4:1 and 0:1. Cells were checked for successful transfection by observing the presence of green signals due to GFP and used for electrophysiological measurements within 48 hours.

### Site-directed Mutagenesis and Generation of Stable Expression Cell Lines of HA-tagged constructs

The full-length hNa_v_1.7 cDNA was available from a former study (19) and was subcloned into the pcDNA5.1/FRT/TO vector using the Gateway PCR Cloning System (ThermoFisher Scientific, Waltham, MA, USA). HA-tags were inserted into the Na_v_1.7 encoding sequence according to the QuikChange protocol (20) using Phusion high-fidelity DNA polymerase and Dpn I restriction endonuclease (New England BioLabs) to generate the non-tagged and HA-tagged hNa_v_1.7 WT or hNa_v_1.7 M899I constructs. The four locations where the HA-tag was inserted are as follows – at the C-terminus (HAC-term), between P148 and P149 in domain 1 S1-S2 (HA1), between L280 and E281 in domain 1 S5-S6 (HA2) and between L293 and E294 in domain 1 S5-S6 (HA3). Oligonucleotides were purchased from Eurofins Genomics. All constructs were verified by restriction pattern analysis and commercial DNA sequencing (Eurofins Genomics).

Inducible stable cell lines expressing the non-tagged and HA-tagged hNa_v_1.7 WT or M899I plasmids were generated using the Flp-In™ T-REx™ kit and the Flp-In™ T-REx™ 293 cell line according to the protocol (Thermo Fischer Scientific, Waltham, MA, USA). Cells were cultured in DMEM 10% FBS, high glucose + L-Glutamin + 15 μg/ml blasticidin and 100 μg/ml hygromycin. Expression was induced by 1 μg/ml doxycyclin and incubating the cells for 24 h at 37°C with 5% CO_2_ before the start of either electrophysiological measurements or immunostaining experiments.

### Whole-cell voltage clamp experiments

Whole-cell voltage clamp of either transiently transfected HEK293t cells or stably expressing HEK293t Flp-in™ cells were performed at room temperature and recorded using the HEKA EPC 10 USB amplifier (HEKA Electronics, Lambrecht, Germany). All recordings were measured in whole-cell configuration and digitally acquired using Patchmaster Next v1.2 (HEKA Electronics, Lambrecht, Germany). The recordings were sampled at a frequency of 100kHz. Leak subtraction was performed digitally using a P/4 protocol following the test pulses. Borosilicate glass capillaries were used to pull recording pipettes using a DMZ puller (Zeitz Instruments GmbH, Martinsried, Germany). Pipette resistances varied between 1 and 3 MΩ. The external bath solution contained (in mM): 140 NaCl, 3 KCl, 1 MgCl_2_, 1 CaCl_2_, 10 HEPES and 20 Glucose with osmolarity maintained around 310mOsm with pH adjusted to 7.3 using CsOH. The internal pipette solution contained (in mM): 10 NaCl, 140 CsF, 1 EGTA, 10 HEPES and 18 Sucrose with osmolarity maintained around 298mOsm with pH adjusted to 7.4 using NaOH. For all cells recorded and considered for further analyses, the series resistance was compensated by at least 70% using series compensation. This ensured that the voltage error did not exceed 8 mV.

After reaching whole-cell configuration, a set of repetitive -10mV depolarizing steps was applied for a duration of three minutes to allow the currents in the cell to stabilize and reach steady-state. After this “monitor” protocol, voltage-dependence of activation was investigated using an activation protocol. This protocol uses a set of 40ms depolarizing test pulses ranging from -90mV to +40mV with a holding potential of - 120mV. The test pulses increase in steps of 10mV, and the inter-sweep interval was maintained at 5s. The peak current (I_peak_) recorded at each applied voltage (V) was used to then calculate the conductance (G) using Eq. 1:

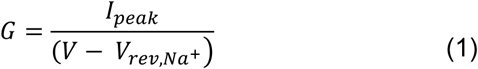

where V_rev,Na+_ is the reversal potential for sodium ions calculated by linear regression fitting of the last three to five voltage sweeps. From the conductances at each voltage step, the maximal conductance (G_max_) is then used to normalize the conductance, which can then be fit to the applied voltage (V) using the Boltzmann function described by Eq. 2:

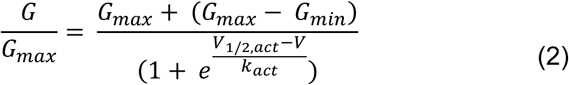

where G_min_ is the minimal conductance, V_1/2,act_ and k_act_ are the voltage at half-maximal channel activation and slope factor respectively. The current densities of the cell can also be calculated using I_peak_ by normalizing the values to the capacitance of the cell. The maximal current densities are then obtained by using the absolute maximal value from the current densities calculated across V. Time to peak was measured as the time taken to reach I_peak_ from the start of the depolarizing step for V between -50mV and +40mV. The onset of fast inactivation was measured by fitting the inactivating phase of the recordings to a single exponential function using Eq. 3:

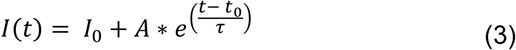

where I(t) is the current amplitude as a function of time t, I0 is the current at t=0, A is the amplitude coefficient and τ is the decay time constant. τ is then plotted against V between -30mV and +40mV.

The voltage-dependence of steady-state fast inactivation was measured using a two-pulse protocol. The sweep starts from a holding potential of -120mV with a 500ms pre-pulse with voltages varying from -140mV to -10mV. This conditions the channels to enter the fast inactivated state, while avoiding entry into slow inactivation (21). This is followed by a 40ms test pulse to a depolarizing voltage of +10mV, to test how many channels are still in the fast inactivated state. The pre-pulses are increased in 10mV steps, with a 10s inter-sweep interval. The peak currents (I_peak_) normalized to the maximal peak current across the applied pre-pulse voltage (V) can be related to V using the Boltzmann function described by Eq. 4:

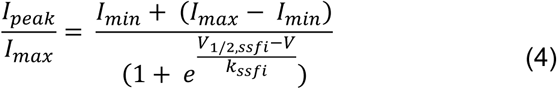

where I_min_ is the minimal peak current across V, V_1/2,ssfi_ and k_ssfi_ are the voltage at half-maximal channel fast inactivation at steady-state and the slope factor respectively.

### Patch clamp data statistics

The raw data recorded were extracted using FitMaster (HEKA Electronics, Lambrecht, Germany) and imported into Igor Pro (WaveMetrics, Portland, OR, USA) to obtain the different gating parameters using in-house scripts. Boltzmann fitting of the normalized conductances and currents to obtain V_1/2,act_, k_act_, V_1/2,ssfi_ and k_ssfi_ were done using GraphPad Prism 5. The final plotting of the data, generation of images and statistical testing was done using GraphPad Prism 5. For statistical testing, a significance level of 95% was used with either the one-way ANOVA with post-hoc Bonferroni corrections or the Kruskal-Wallis with post-hoc Dunn’s corrections used based on the normality of the data. Error bars typically represent the 95% confidence interval of the mean unless stated otherwise. Although ANOVA was run, only the significant differences from WT are shown in figures and tables (* p<0.05). Generation of all figures in this study were done using Inkscape.

### Immunocytochemistry and confocal imaging

For immunostaining of the stable HEK293 Flp-in™ cell lines expressing either non-tagged and HA-tagged hNa_v_1.7 WT/M899I, cells were seeded onto coverslips and incubated with 1µg/µL doxycycline for ∼24h before staining. Post 24h, coverslips were incubated with 4% PFA at room temperature for 15 minutes and washed with PBS thrice to remove excess PFA, Permeabilization of the membrane was done using 0.1% Triton X-100. The permeabilization step wasskipped for the extracellularly located tags. Blocking solution made of PBS and 5% normal goat serum were added and incubated at room temperature for 60 minutes. 200µL of primary antibodies targeting the HA tag (Cell Signalling Technologies, #3724S) diluted in blocking solution was added before being incubated overnight in a wet chamber at 2-8°C. The next day, coverslips were washed three times with PBS to remove excess unbound primary antibodies. Secondary antibodies conjugated with a fluorophore (Life Technologies, #A-21429, Alexa Flour 555) were then added and incubated for 60 minutes at room temperature without exposure to light. Three washings with PBS were performed to remove unbound secondary antibodies. DAPI was added to the second wash (NucBlue Fixed Cell ReadyProbes Reagenz, Life Technologies, #R37606) to stain the nuclei. The slides were finally mounted with 10µL of fluorescent mounting medium (Dako) and stored at 4°C in the dark for imaging at a later timepoint.

For immunostaining with antibodies targeting Na_v_1.7, HEK293t cells seeded into coverslips were transiently transfected with hNa_v_1.7 WT or hNa_v_1.7 M899I. Cells were then checked for GFP expression to validate successful transfection. The coverslips were then fixed by incubation with 4% PFA at room temperature for 15 minutes, with addition of 0.1% Triton X-100 to permeabilize the cells. To allow the coverslips to be used at a later time point, coverslips were also incubated with 0.05% sodium azide (diluted in PBS) and sealed with parafilm onto a slide plate. These cells were then exposed to primary antibodies targeting Na_v_1.7 (Abcam, #ab85015) and incubated overnight in a wet chamber at 2-8°C. The next day, secondary antibodies (Life Technologies, #A21434, Alexa Flour 555) conjugated with flurophores that target the primary antibody were added and incubated for 2h before using for confocal imaging. DAPI (NucBlue Fixed Cell ReadyProbes Reagenz, Life Technologies, #R37606) was used to stain the nuclei of the cells.

Confocal imaging of the immunostained cells was performed using a LSM700 confocal microscope (Zeiss, Germany) with a Plan-Apochromat 63x/1.4 NA oil objective (Zeiss, Germany) and acquired digitally using the ZEN black software (Zeiss, Germany). The intensities of the laser was kept constant throughout all images acquired to allow for accurately observing qualitative differences between the images. Post-processing of the images including the addition of scale bars, merging of the channels and adjustment of the brightness of the DAPI signal were done using ImageJ. The DAPI signal brightness was artificially enhanced in the non-permeabilized coverslips where necessary to allow for the nuclei to be located in the images.

### Multiple sequence alignment

Protein sequences for the nine hNa_v_ subtypes (hNa_v_1.1-1.9), the Na_v_1.7 of the rat (rNa_v_1.7), mouse (mNa_v_1.7) and rabbit (rabbitNa_v_1.7), the Na_v_1.4 of the electric eel (eeNa_v_1.4) and the Na_v_ of the American cockroach (Na_v_PaS) were obtained from the Uniprot database (22) (Accession codes - hNa_v_1.1 Q8NEY1, hNa_v_1.2 Q99250, hNa_v_1.3 Q9NY46, hNa_v_1.4 P35499, hNa_v_1.5 Q14524, hNa_v_1.6 Q9UQD0, hNa_v_1.7 Q15858, hNa_v_1.8 Q9Y5Y9, hNa_v_1.9 Q9UI33, rNa_v_1.7 O08562, mNa_v_1.7 Q62205, rabbitNa_v_1.7 Q286444, eeNa_v_1.4 P02719 and Na_v_PaS D0E0C2). Sequences were loaded into Jalview and multiple sequence alignment was performed on these sequences using the T-Coffee package provided by JABAWS (23). The sequences were truncated to contain only the region around the hNa_v_1.7 M899I residue and shaded according to the level of sequence conservation using Boxshade. The black shades represent fully conserved residues, grey shades represent partially conserved residues, and no shade represents non-conserved residues.

### Homology Modelling and molecular visualization

Homology modelling of the hNa_v_1.7 WT, M899A, M899C, M899F, M899I, M899L and M899V was done using either Modeller v9.23 or v10.2 (24) with the cryo-EM structure of the hNa_v_1.7 in a fast inactivated state used as the template (PDB ID:6J8G) (25). The structures do not contain the N- and C-termini and the intracellular inter-domain linkers D1-D2 and D2-D3. The structures were first prepared by removing the ß subunits and other heteroatoms (such as detergents, lipids, ions etc). For introducing the variants, M899 in the sequence alignment was changed to the amino acid of interest. To introduce the various extracellular HA-tagged structures (HA1, HA2 and HA3), the HA sequence was added in-between the two residues where the HA-tag must be placed before alignment to the template. For all models missing atoms between domain 2 S2 and domain 2 S3 (amino acids 815-818) were filled using Modeller. A total of 100 models were generated for each variant. For the HA-tagged structures, a total of 15 models were generated. The best model was chosen based on the most negative Discrete Optimized Protein Energy Score (DOPE-HR). The best model was chosen for further visualization or usage in coarse-grained molecular dynamics (MD) simulations. Molecular visualization and image generation was done using ChimeraX (26).

### Coarse-grained molecular dynamics simulations

The best models of the WT or M899 variants were used for coarse-grained molecular dynamics (MD). Coarse-grained MD was performed using GROMACS v2022/v2022.3 (27, 28) with the Martini v3 forcefield (29). Briefly, the best model was first aligned to the membrane bilayer using the PPM webserver (30). The aligned structure was then used to generate the coarse-grained structure and topology using Martinize2 (31). To maintain the secondary and tertiary structural information, elastic networks with a force constant of 700 kJ/mol/nm^2^ were added between the non-bonded backbone beads who are within a distance between 0.3nm to 0.9nm. Side-chain dihedral angle corrections were introduced using scfix (32). The coarse-grained structure was then embedded in a pure (100%) polyphosphatidylcholine (POPC) membrane with water and 150mM NaCl using Insane (33). The total charge of the system was kept neutral, and the system itself was contained in a 18×18×18nm^3^ cubic box with at least 2.5nm between two periodic images of the protein. The system underwent two minimization steps using the steepest descent algorithm – first with a maximal force cut-off of 1000 kJ mol^-1^ nm^-1^ and the second with a reduced value of 500 kJ mol^-1^ nm^-1^. The minimized system was then equilibrated shortly for 2ns in a NVT ensemble in 2fs timesteps to allow for the stabilization of the box size and volume. Post short NVT-equilibration, a longer equilibration was performed in a NVT ensemble for 200ns in 20fs time steps and another longer equilibration in a NPT ensemble for 50ns in 20fs time steps. At the end of the equilibration steps, temperature and pressure were checked to ensure it remained stable at 310K and 1 bar respectively. Temperature was maintained using a v-rescale thermostat while pressure was maintained using the Parrinello-Rahman barostat in a semi-isotropic manner. For all equilibration steps a backbone restraint of 1000kJ/mol was maintained. Post equilibration, the system underwent an unbiased and unrestrained production run using the md integrator for a total of 4µs in 20fs steps. For each system, three 4µs replicas were generated.

The trajectories of each replica were downsampled to intervals of 10ns and corrected for periodic boundary conditions and roto-translation of the system. The protein was then isolated for further analyses. Root mean square deviation and fluctuations (RMSD and RMSF respectively) of the backbone beads were calculated using the ‘rms’ and ‘rmsf’ modules of GROMACS. The first 1µs of the trajectories were then discarded from further analyses, resulting in 3µs per replica and a total of 9µs per system. The trajectories were then split into individual frames, and the pore radius of each frame was calculated using HOLE 2.0 (34). The center of geometry of each individual frame was used as the starting point, with the radius calculated along the z vector. The end radius was set to 15A°, and the radius file was modified to include the radii of the beads. Pore radii were calculated for every frame in steps of 0.25A° along the z vector. The average pore radius was calculated for each pore coordinate as the average value over the 903 frames for each system. Pore radius values were only plotted between pore coordinate values normalized to the center of geometry between -16A° intracellularly and 15A° extracellularly. Volumes were calculated as the sum of the volumes of each individual slice approximated by π * (radius at slice)^2^ * h, where h is 0.25A°. Volumes were always calculated per frame, and the volumes over the 903 frames were plotted as a violin plot. The helix bending angles of the S5, P1, P2 and S6 helices for every trajectory were obtained using Bendix (35). An average value per residue of the helix was calculated for every protein system by averaging over the three replicas.

Kullback-Leibler divergence was used to measure the differences in the probability densities of either pore radii or helix bending angles of WT (P(x)) compared to a variant (Q(x)) using Eq. 5:

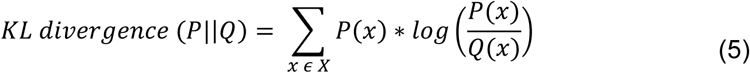

where X represents the sample space to which P and Q belong.

The extraction of individual frames, calculation of the probability densities and generation of the final graphs and violin plots were all done using in-house custom scripts in Python3.

## Results

### hNa_v_1.7 M899 is highly conserved and located in the pore module of domain 2

The hNa_v_1.7 M899 residue is situated in the pore module of domain 2, more specifically in the re-entrant pore loop between the S5 and P1 helices (Figure 1A). Multiple sequence alignment revealed that the residue is fully conserved across not only human Na_v_s such as the cardiac Na_v_1.5, but also Na_v_1.7 of various mammalian species and the Na_v_1.4 of the electric eel (Figure 1B). The Na_v_ of the American cockroach (Na_v_PaS) was instead found to have a phenylalanine (Figure 1B). Although not directly in the permeation pathway, the residue is in proximity to the selectivity filter motif “DEKA” (Figure 1C) and tightly packed in a hydrophobic pocket (Figure 1D). The residue M899 is also close to the semi-conserved cysteine residue involved in the binding of the ß2 subunit (25) (Figure 1D, C884).

### hNa_v_1.7 M899I causes complete loss of function in HEK293T cells

The hNa_v_1.7 M899I and hNa_v_1.7 WT were transiently transfected into HEK293t cells and co-transfected with GFP as a reporter gene, and characterized using whole-cell voltage clamp. While hNa_v_1.7 WT showed robust inward sodium currents with fast inactivation kinetics, the hNa_v_1.7 M899I did not show any observable currents (Figure 2A and Table 1). The current densities of hNa_v_1.7 M899I were indistinguishable from HEK293t cells transfected with GFP only (*p<0.05; Figure 2B-C and Table 1). The loss of function caused by M899I could not be rescued by lowering incubation temperatures or by co-expression of ß2 (*p<0.05; Figure 2D and Table 1), as observed for some trafficking-deficient loss-of-function Na_v_ variants (37). The loss of function caused by mutating M899 was not limited to Na_v_1.7. Due to the fully conserved nature of this residue, mutating the equivalent residue in hNa_v_1.2 or hNa_v_1.5 also resulted in a complete loss of function when transfected in HEK293t cells (*p<0.05; Figure 2E and Table 1).

**Figure 2:**
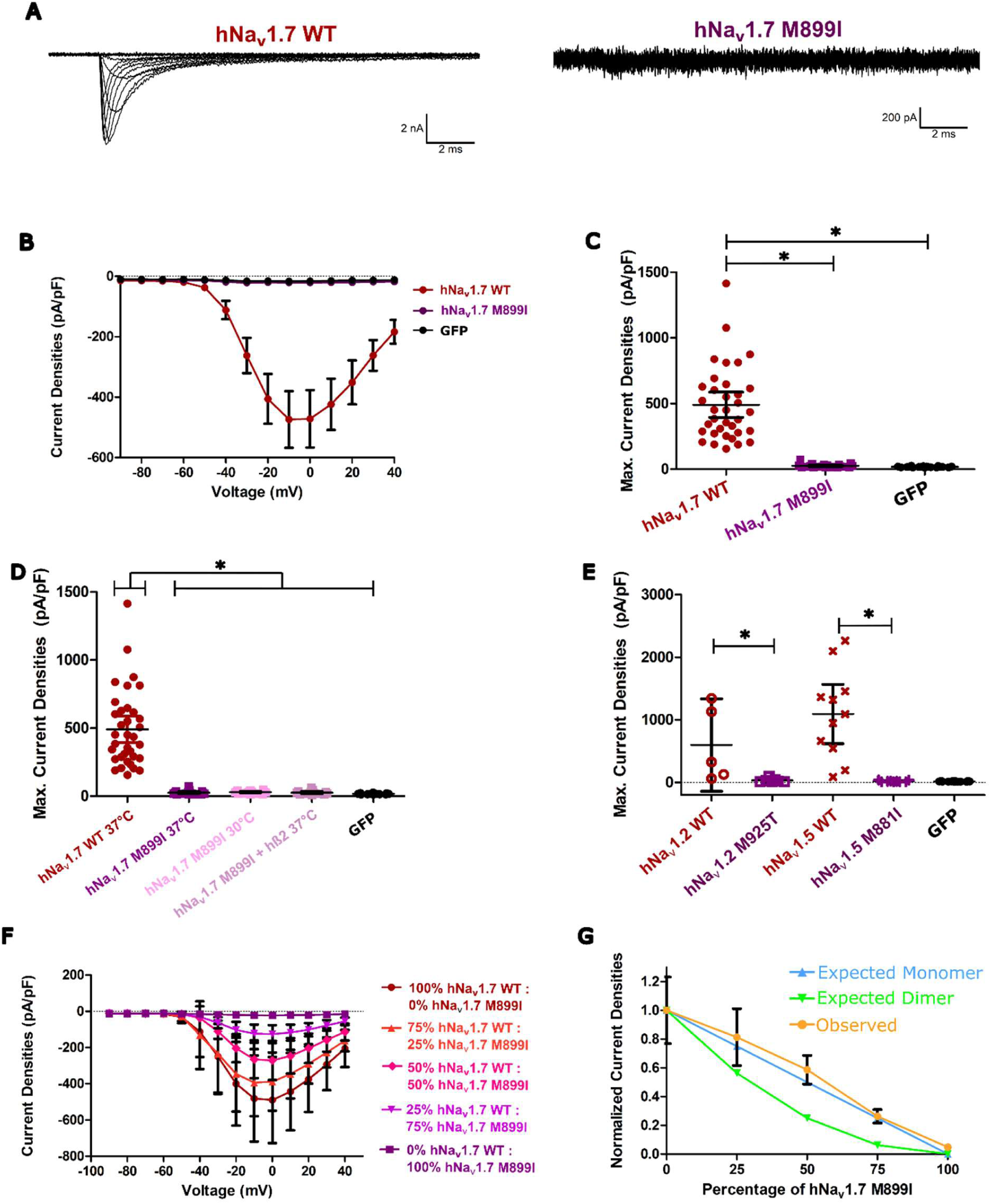
hNa_v_1.7 M899I causes complete loss-of-function in HEK293t cells. **A)** Representative traces of the hNa_v_1.7 WT (left) and hNa_v_1.7 M899I (right) obtained by voltage-clamp of transiently transfected HEK293t cells. While hNa_v_1.7 WT shows robust inward sodium currents hNa_v_1.7 M899I shows visually unobservable currents across all voltage steps applied. **B)** Mean current densities vs applied voltage steps. hNa_v_1.7 M899I is indistinguishable from HEK293t cells transfected only with GFP (negative control) **C)** Dot plot of the maximal current densities. The maximal current densities for hNa_v_1.7 M899I are indistinguishable from the negative control (GFP). **D)** Dot plot of the maximal current densities of hNa_v_1.7 WT incubated at 37°C, hNa_v_1.7 M899I incubated at 37 or 30°C, hNa_v_1.7 M899I co-transfected with the ß2 subunit and incubated at 37°C and the negative control (GFP). Neither decreasing the incubation temperature nor adding secondary proteins like the ß2 subunit rescued the current densities of hNa_v_1.7 M899I. **E)** Maximal current densities of hNa_v_1.2 or hNa_v_1.5 WT and the mutated residues corresponding to M899 in hNa_v_1.7 (M925T in hNa_v_1.2 and M881I in hNa_v_1.5). Mutation of the M899-corresponding hNa_v_1.2 or hNa_v_1.5 residues also result in complete loss of function when transfected in HEK293t cells. **F)** The mean current densities vs applied voltage (protocol in A inset) with various ratios of hNa_v_1.7 WT and hNa_v_1.7 M899I transiently co-transfected in HEK293t cells. The current densities show a trend of decreasing as the amount of hNa_v_1.7 M899I increases. **G)** Plot of the normalized current densities against the percentage of hNa_v_1.7 M899I present in the various ratios. The observed trend follows very closely the ideal relationship for monomeric existence of the channels, providing no evidence for dominant negative effects. Dominant negative effects are typically considered plausible, when the presence of 50% of non-functional channels result in a decrease of ∼75% of the current densities (3). Error bars represent the 95% confidence interval of mean except in panel G, where it represents the standard error of the mean. * p<0.05

**Table 1.**
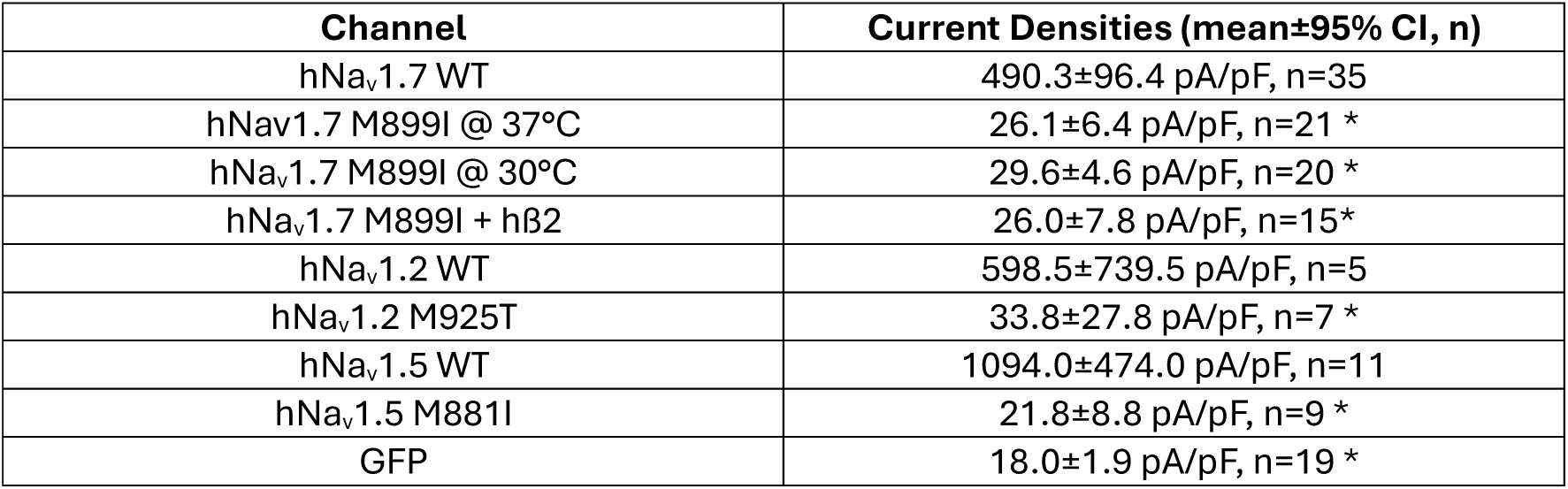
Current densities of HEK293t cells transiently transfected with either hNa_v_1.7 WT, hNa_v_1.7 M899I at incubation temperatures of either 37°C or 30°C, hNa_v_1.7 M899I co-transfected with the hß2 subunit, hNa_v_1.2 WT, hNa_v_1.2 M925T, hNa_v_1.5, hNa_v_1.5 M881I or only reporter protein GFP as a control. Values are always written as the mean ± the 95% confidence interval (95% CI) of the mean, number of cells/data points. * p<0.05 when compared to hNa_v_1.7 WT.

| Channel | Current Densities (mean±95% CI, n) |
| --- | --- |
| hNav1.7 WT | 490.3±96.4 pA/pF, n=35 |
| hNav1.7 M899I @ 37°C | 26.1±6.4 pA/pF, n=21 * |
| hNav1.7 M899I @ 30°C | 29.6±4.6 pA/pF, n=20 * |
| hNav1.7 M899I + hβ2 | 26.0±7.8 pA/pF, n=15* |
| hNav1.2 WT | 598.5±739.5 pA/pF, n=5 |
| hNav1.2 M925T | 33.8±27.8 pA/pF, n=7 * |
| hNav1.5 WT | 1094.0±474.0 pA/pF, n=11 |
| hNav1.5 M881I | 21.8±8.8 pA/pF, n=9 * |
| GFP | 18.0±1.9 pA/pF, n=19 * |

Heterozygous variants were shown to influence the gating of WT Na_v_s due to the dimerization of the alpha subunits (5). The here presented patient was shown to carry the M899I variant heterologously, with no variants reported in the SCN9A of the other allele (17). It could therefore be possible that the variant may interact with the Na_v_1.7 encoded by the other allele to cause dominant negative effects. To test this possibility, varying amounts of hNa_v_1.7 WT and M899I were co-transfected in HEK293t cells and current densities in relation to the amount of M899I were quantified as previously described (3).

The current densities showed a decrease as the amount of co-transfected non-functional M899I increases (Figure 2F and Table 2). Plotting the normalized current densities of the various ratios as a function of the amount of non-functional M899I showed that this decrease was linear (Figure 2G, Table 2). This behaviour is expected if the two channels would remain monomeric or if the dimerization would not lead to a dominant negative effect of the variant (Figure 2G and Table 2).

**Table 2.**
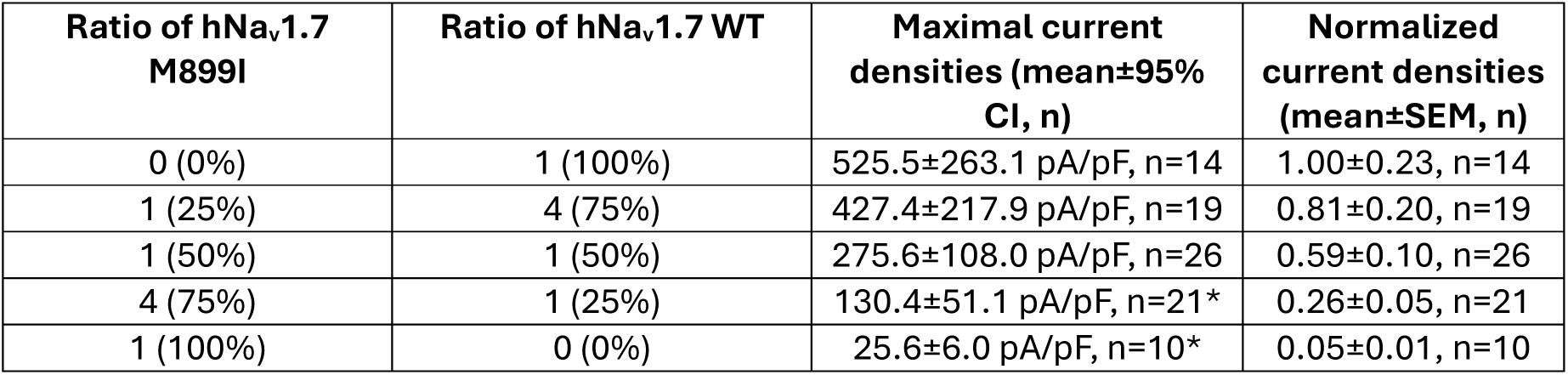
Variation of normalized current densities when varying the quantity of hNa_v_1.7 M899I co-transfected transiently with hNa_v_1.7 WT in HEK293t cells. Quantities of plasmids are normalized as ratios. Normalized current densities are obtained by normalizing the maximal current densities to the average current density of the control 0:1 ratio (0% hNa_v_1.7 M899I and 100% hNa_v_1.7 WT). Data is represented as mean±95% confidence interval (95% CI) of the mean for current densities and as mean± standard error of mean (SEM) for the normalized current densities.n represents the number of cells/data points. * p<0.05 when compared with control 0:1 ratio (0% hNa_v_1.7 M899I and 100% hNa_v_1.7 WT).

| Ratio of hNav1.7 M899I | Ratio of hNav1.7 WT | Maximal current densities (mean±95% CI, n) | Normalized current densities (mean±SEM, n) |
| --- | --- | --- | --- |
| 0 (0%) | 1 (100%) | 525.5±263.1 pA/pF, n=14 | 1.00±0.23, n=14 |
| 1 (25%) | 4 (75%) | 427.4±217.9 pA/pF, n=19 | 0.81±0.20, n=19 |
| 1 (50%) | 1 (50%) | 275.6±108.0 pA/pF, n=26 | 0.59±0.10, n=26 |
| 4 (75%) | 1 (25%) | 130.4±51.1 pA/pF, n=21* | 0.26±0.05, n=21 |
| 1 (100%) | 0 (0%) | 25.6±6.0 pA/pF, n=10* | 0.05±0.01, n=10 |

### hNa_v_1.7 M899I is expressed in the cell membrane

The indistinguishable current densities of hNav1.7 M899I and GFP-transfected HEK293T control cells prompted us to examine whether the variant was expressed at the plasma membrane. Confocal imaging of HEK293t cells transfected with WT or M899I and incubated with antibodies targeting Na_v_1.7 showed clear fluorescent signals throughout the cell body indicating the presence of Na_v_1.7 WT or M899I in these cells (Figure S1).

To ensure localization of the mutated Na_v_1.7 in the cell membrane and not merely intracellularly, extracellular tagging strategies were applied. An HA-tag was inserted into one of three extracellular locations in hNa_v_1.7 (HA1, HA2 and HA3; Figure 3A). In immunostainings against the HA-tag, non-tagged channels did not show any signals (Figure 3B). We also inserted an HA-tag into the C-terminus of hNa_v_1.7, to test for antibody specificity. The intracellular tag can be detected only following permeabilization of the membrane (Fig 3C). Confocal imaging of HEK293t cells stably expressing hNa_v_1.7 WT or M899I channels with HA-tags inserted in either of three locations was performed in a non-permeabilized manner with antibodies targeting the HA-tag. Both WT and M899I HA-tagged channels showed a clear signal irrespective of the position of the HA tag, visible as a red rim around the cell membrane (Figure 3D, white arrows). Whole-cell patch clamp of HEK293t cells stably expressing HA-tagged hNa_v_1.7 WT showed minimal to no change in both expression and gating properties irrespective of the location of the HA-tag when compared to non-tagged channels (Figure S2 A-J and Table S1). These results show that hNa_v_1.7 M899I channels are expressed in the membrane.

**Figure 3:**
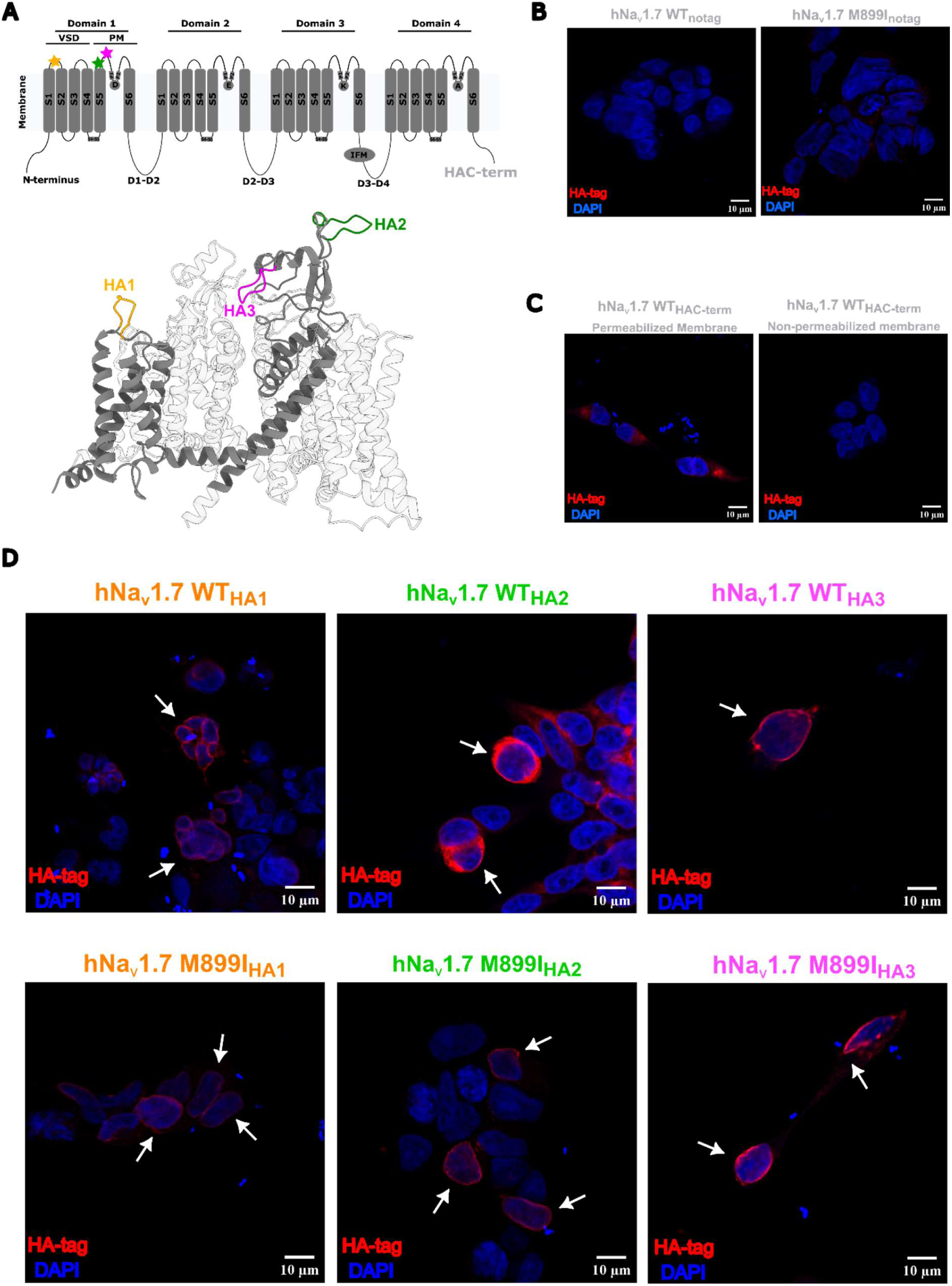
hNa_v_1.7 M899I is expressed in the membrane. **A)** 2-D schematic of Na_v_1.7 with extracellular HA-tag positions marked by colored stars and the C-terminal HA tag represented by HA-Cterm. Below is the 3-D representation of the hNa_v_1.7 cryo-EM structure in fast inactivated state (PDB ID: 6J8G) (36) with three different extracellular HA-tag positions (HA1, HA2 and HA3) in domain 1 via homology modelling using Modeller v10.2 (24). Domain 1 is shown in opaque grey while the other parts of the channel are made transparent for clarity. **B)** Confocal imaging of HEK293t cells stably expressing hNav1.7 WT or M899I without HA-tag. No HA-signal can be observed in either WT or M899I. **C)** Confocal imaging of HEK293t cells transfected with intracellularly HA-tagged hNav1.7 WT and immunostained with antibodies targeting the HA tag. Staining was performed either with (left) or without (right) permeabilization of the membrane. An HA-signal can only be observed when the membrane is permeabilized to allow the antibodies access to the intracellular HA-tag. **D)** Confocal imaging of HEK293t cells stably expressing hNa_v_1.7 WT or M899I with HA tag in either of the three different locations (HA1, HA2 or HA3, red). The top row represents WT channels while the bottom row represents M899I channels. The first column represents channels without any tags attached, while each of the next three columns represent in order HA1, HA2 and HA3HA-tagged WT or M899I channels show a clear rim-like HA staining around the boundary of the cell (white arrows) irrespective of the HA-tag location. The scale bar is shown in the bottom right corner for all confocal images.

### hNa_v_1.7 M899I induces outer-pore collapse without altering local helix bending in CGMD simulations

The hNa_v_1.7 M899I variant is expressed in the membrane but does not produce currents. Thus, we turned to *in-silico* modelling to understand the impact of the variant on channel permeation. To elucidate on the structural defects induced by hNa_v_1.7 M899I, CGMD using Martini3 was used to simulate homology models of hNa_v_1.7 WT and M899I. The channels were embedded into a pure POPC membrane surrounded by water and ions.

To characterize whether the amino acid substitutions altered local geometry around the packed region of residue M899, the bending angles of the helices that make up the PM of domain 2 (S5, P1, P2 and S6) were calculated using Bendix (35). The angles were calculated as an average of the three replicas, and the KL divergence between the probability densities of hNa_v_1.7 WT and M899I for each pore helix was calculated using Eq. 5. Irrespective of the pore helix, hNa_v_1.7 WT and M899I did not show major differences in the bending angles (Figure S3 A-D). The average root mean square deviations (RMSD) and fluctuations (RMSF) between hNa_v_1.7 WT and M899I were very similar (Figure S3 and S4).

The pore radius was calculated on the post-processed frames of the last 3µs of each replica and investigated for any changes. The average pore radius showed a large reduction at the outer pore near the outer vestibule (OV) region, which was confirmed to be due to an inherent difference in the probability densities of the pore radii across the 903 frames between hNa_v_1.7 WT and M899I, as shown by the large KL-divergence value corresponding to the bottlenecked region (Figure 4A, upper black arrow). A secondary local maximum could be observed in the KL-divergence plot corresponding to the center-of-geometry of the structure (Figure 4A, lower black arrow). Visualization of the average 3-D pore profile between hNa_v_1.7 WT and M899I confirmed the 2-D pore radius plots, showing a constriction at the outer pore in the region between the selectivity filter (SF) and the OV (Figure 4B). To better quantify these parameters, violin plots of the pore radii at the OV (corresponding to the highest KL-divergence value), the volume between SF and OV (corresponding to the bottleneck in the 3D-pore profile), the radii at the center-of-geometry (CoG) of the protein backbones (corresponding to the second local maxima in the KL-divergence plot) and the central cavity volume (quantifiable parameter close to the CoG) were graphed. The average pore radius at the OV was drastically reduced in hNa_v_1.7 M899I, taking up more values below 1A° compared to hNa_v_1.7 WT (Figure 4C). This also corresponded with a similar reduction in the volume between the SF and OV, with average value in hNa_v_1.7 M899I around 50% lesser than hNa_v_1.7 WT (Figure 4D). The radius at CoG and the central cavity volume do not show major differences in the average values, although hNa_v_1.7 M899I was observed to have a longer tail towards smaller values compared to hNa_v_1.7 WT (Figure 4E and F).

**Figure 4:**
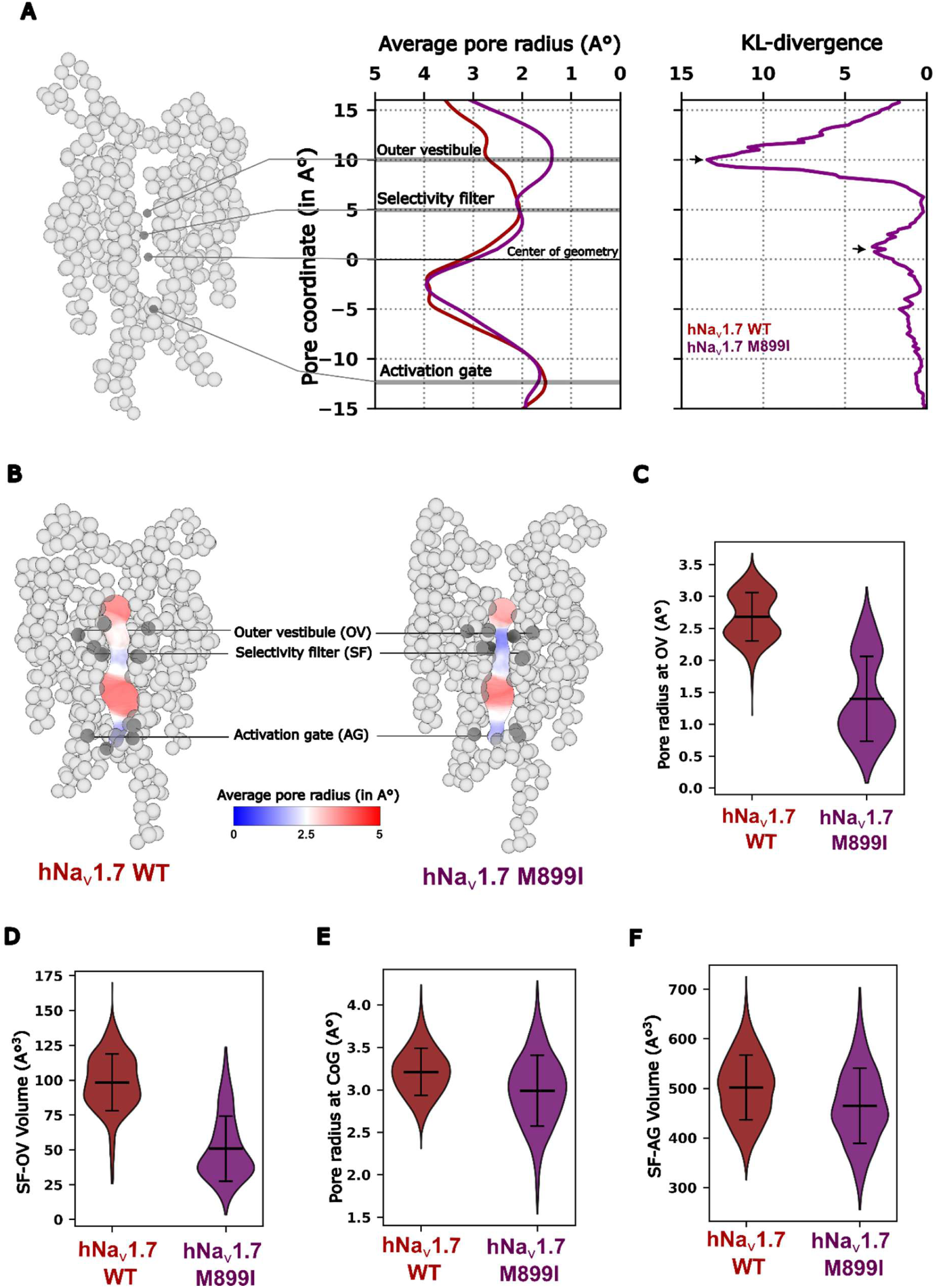
hNa_v_1.7 M899I causes a bottlenecking of the outer pore. **A)** Average pore radius and the corresponding KL-divergence vs pore coordinate between hNa_v_1.7 WT and M899I. The pore module of the 3D structure is shown for reference to highlight important regions along the pore. The average pore radius is lowered in hNa_v_1.7 M899I at the outer pore region, which also shows a large KL-divergence value (first black arrow). A second, smaller KL-divergence value is observed close to the center-of-geometry of the structure (second black arrow). **B)** Average 3-D pore profiles of hNa_v_1.7 WT and M899I. The pore profile is colored based on the the average pore radius, ranging from 0A° (blue) to 5A° (red). The three important regions along the pore – outer vestibule (OV), selectivity filter (SF) and activation gate (AG) – are highlighted. A bottlenecking can be observed in the region between the OV and SF in hNa_v_1.7 M899I. Violin plots of **C)** pore radius at OV, **D)** the volume of the pathway between SF and OV, **E)** pore radius at the center-of-geometry (CoG) of the protein backbones and **F)** the central cavity volume approximated as the volume of the pathway between SF and AG. The pore radius at OV and the SF-OV volume is drastically reduced in M899I, with minor differences in distribution tails observed in the pore radius at CoG and the central cavity volume. Error bars represent the standard deviation.

### Graded constriction of the outer pore by hydrophobic substitutions at M899

To assess the sensitivity of the position 899 for amino acid substitutions and the subsequent effect on pore geometry, we chose the following hydrophobic amino acids other than isoleucine: cysteine due to its smaller size but similar chemical group compared to methionine, alanine as the smallest hydrophobic amino acid, phenylalanine as a bulky aromatic amino acid, and valine and leucine as the other branched-chain amino acids. Models were made for each of these amino acids and simulated in a similar manner to hNa_v_1.7 M899I. The average and per replica RMSD and RMSF were very similar (Figure S3 and S4).

The average pore radius differed among the M899 substitutions, revealing varying degrees of outer-pore bottlenecking. hNav1.7 M899V exhibited pore-radius values most similar to those of M899I, whereas M899L and M899F more closely resembled hNa_v_1.7 WT. In contrast, M899A and M899C displayed intermediate levels of constriction (Figure 5A). The KL-divergence captures this graded behaviour more clearly, with peaks observed in a similar order around 10A° (Figure 5A, rightmost panel and upper black arrow). Once again, a second local maximum is observed around 0A°(Figure 5A, rightmost panel and lower black arrow). Quantification of the pore radius and the central cavity volume showed no major differences in the mean values between the various substitutions, with minor alterations in the distribution shape (Figure S6 A-B). The violin plots of the pore radius at the OV show that the mean values follow the same pattern across the hNa_v_1.7 variants – WT > M899L ∼ M899F > M899A∼M899C > M899V > M899I. The SF-OV volume plots also show that this trend is translated in 3-D, with the observation that the average volume of hNa_v_1.7 M899V was smaller than M899I (Figure 5C). However, a closer look at the distribution of the individual values shows that hNa_v_1.7 M899I samples more values that are lower than the mean compared to M899V (Figure 5C, teal arrows).

**Figure 5:**
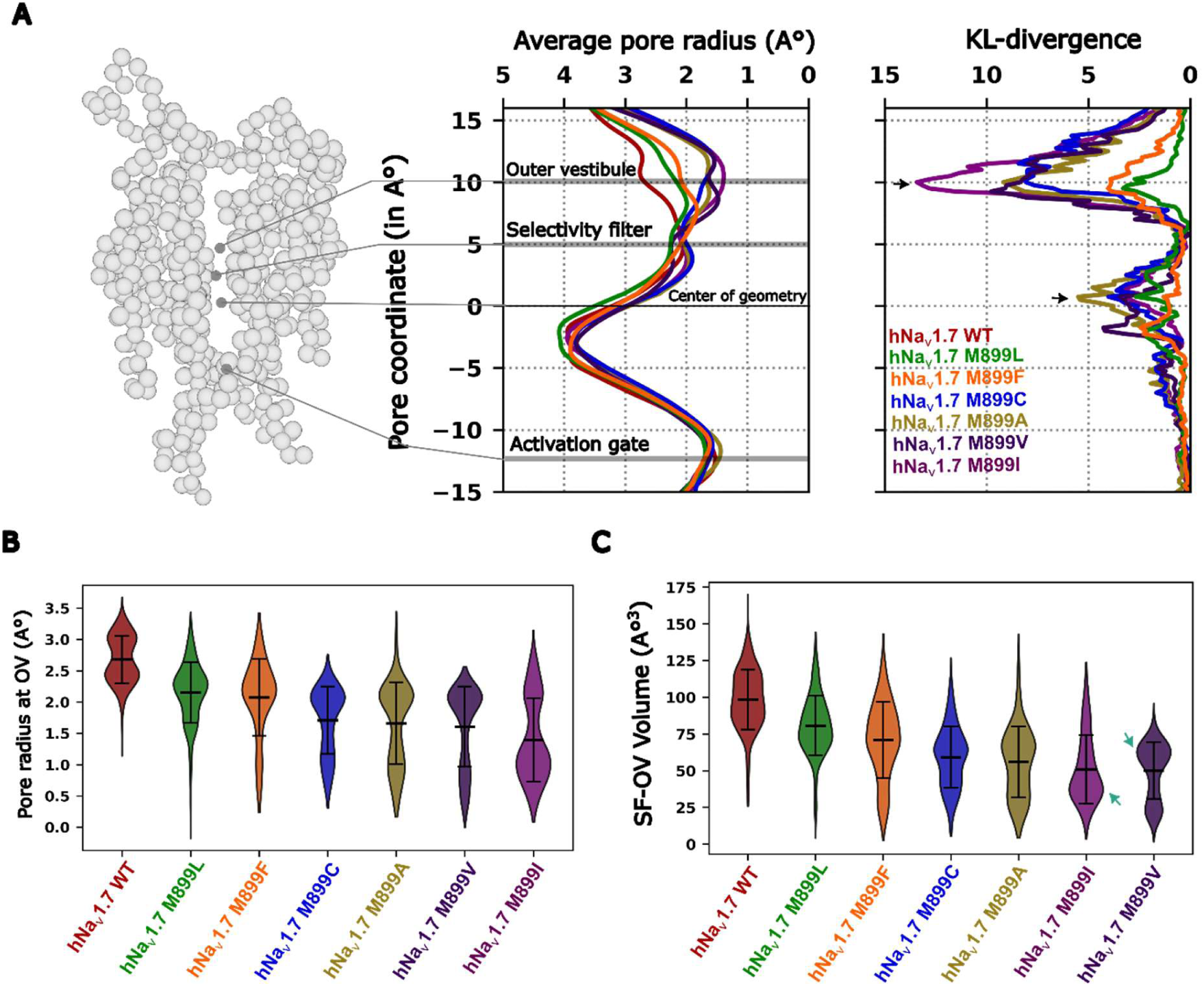
hNa_v_1.7 M899I predicted by coarse-grained molecular dynamics to cause a bottleneck of the outer pore. **A)** Average pore radius and the corresponding KL-divergence vs pore coordinate between hNa_v_1.7 WT and the M899 substituted structures. The pore module of the 3D structure is shown for reference to highlight important regions along the pore. The average pore radius is lowered in a graded fashion at the outer pore region, which also is visible when looking at the large KL-divergence value in the outer pore (first black arrow). A second, smaller KL-divergence value is observed close to the center-of-geometry of the structure (second black arrow). Violin plots of **B)** pore radius at OV and **C)** the volume of the pathway between SF and OV. hNa_v_1.7 M899L and M899F show similar pore radius at the OV compared to hNa_v_1.7 WT, with hNa_v_1.7 M899V showing values closer to M899I and hNa_v_1.7 M899C and M89A in between. When looking at the volume, the trend is mostly the same except for hNa_v_1.7 M899V – M899V shows a lower mean volume compared to M899I. However, a look at the distribution of the individual values shows that hNa_v_1.7 M899I has more values below the mean compared to M899V (teal arrows). Error bars represent the standard deviation.

### *In-vitro* current densities of hNa_v_1.7 M899 substitutions closely follow *in-silico* predictions

The *in-silico* predictions highlighted the graded bottleneck of the outer pore radius and volume of the pathway between SF and OV based on the amino acid substituted for M899. These *in-silico* predictions suggest that the *in-vitro* measurable current densities of the hNa_v_1.7 constructs would follow the increasing bottleneck of the outer pore (from high to low): WT > M899L > M899F > M899C > M899A > M899V >= M899I.

To validate the *in-silico* predictions, we utilized whole-cell patch clamp to measure the gating parameters of the investigated M899 substitutions using transfected HEK293t cells. We were able to replicate the in-silico predictions using quantification of the current densities: hNa_v_1.7 WT has the highest, M899F has the second highest – M899V has the second lowest current densities, and M899I has the lowest current density (Figure 6A and B). hNa_v_1.7 M899L showed the third lowest current densities, with M899C and M899A somewhere in-between (Figure 6A-B and Table 3). Statistically, however, only the branched-chain amino acids - M899I, M899V and M899L – showed significant differences when compared to hNa_v_1.7 WT (p < 0.05; Figure 6B and Table 3).

**Figure 6:**
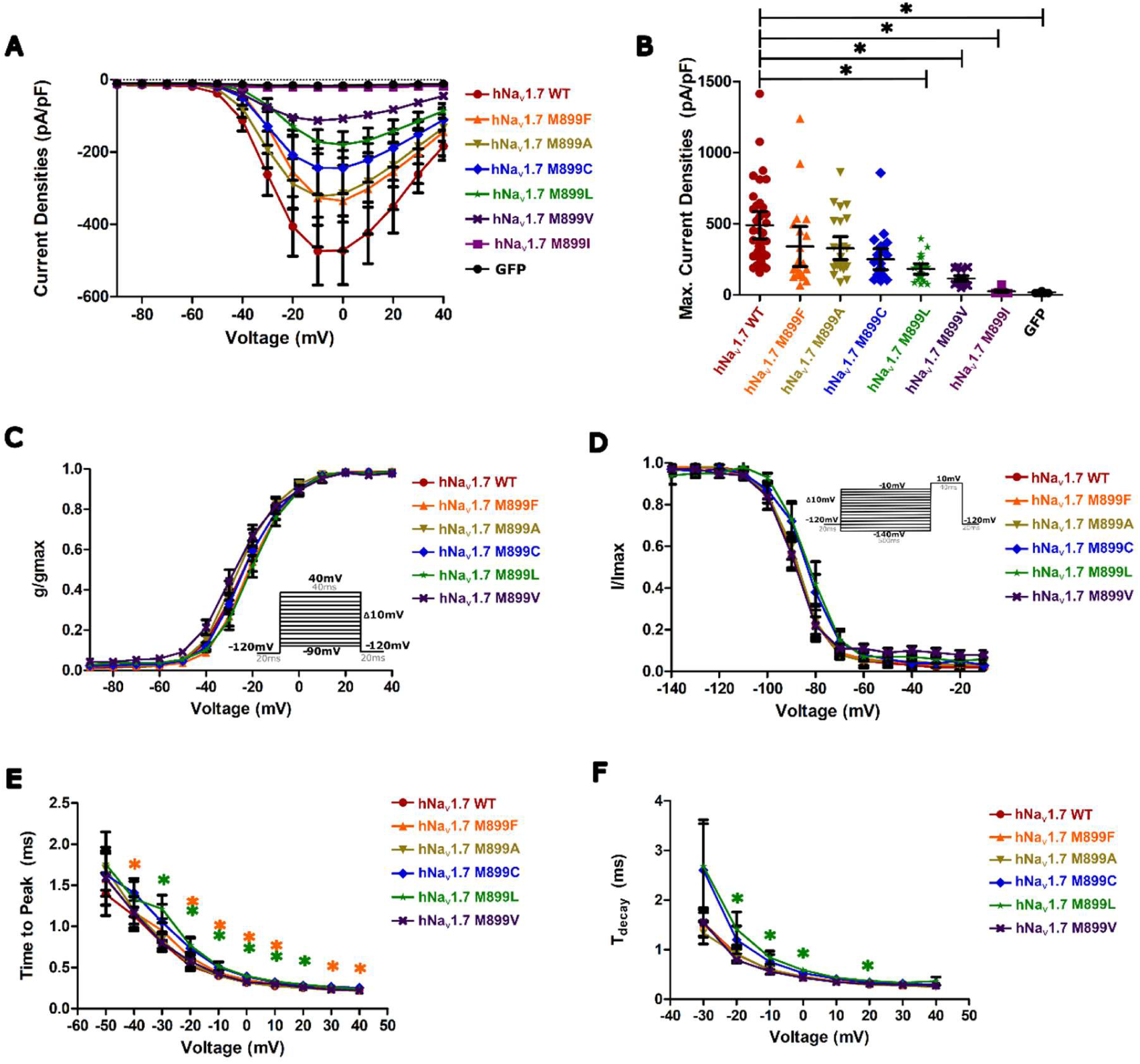
Current densities of hNav1.7 M899 mutations with five amino acids closely follow the predictions made *in silico* with minimal alteration in other gating parameters. **A)** Mean current densities vs applied voltage steps (protocol in C inset) of HEK293t transfected with either hNa_v_1.7 WT, only GFP or either of the six substitutions of M899 (M899F, M899A, M899C, M899L, M899V or M899I). . **B)** Dot plot of the maximal current densities of HEK293t transfected with either hNa_v_1.7 WT, only GFP as a negative control or either of the six substitutions of M899 (M899F, M899A, M899C, M899L, M899V or M899I).. All variants show channel expression, with M899V and M899L significantly lower than WT. *p<0.05. **C)** Normalized conductance curves measured using the protocol in inset for HEK293t transfected with either hNa_v_1.7 WT or either of the five substitutions of M899 (M899F, M899A, M899C, M899L or M899V). **D)** Normalized current curves measured using the protocol in inset, for HEK293t transfected with either hNa_v_1.7 WT or either of the five substitutions of M899 (M899F, M899A, M899C, M899L or M899V). **(E)** Time to peak and **(F)** onset of fast inactivation kinetics for HEK293t transfected with either hNa_v_1.7 WT or either of the five substitutions of M899 (M899F, M899A, M899C, M899L or M899V).. The error bars represent the 95% confidence interval of the mean. * p<0.05. Color of * represents the variant significantly different from WT.

**Table 3.** Current densities and gating parameters of HEK293t cells transiently transfected with either hNa_v_1.7 WT or various substitutions of the M899 residue (M899I, M899A, M899C, M899F, M899L or M899V). GFP transfection acts as negative control. Gating parameters include the V_1/2,act_, slope_act_, V_1/2,ssfi_ and slope_ssfi_. Data are always presented as mean±95% confidence interval (95% CI) of the mean. n represents the number of cells/data points. *p <0.05 when compared to hNa_v_1.7 WT.

|  | hNav1.7 WT | hNav1.7 M899A | hNav1.7 M899C | hNav1.7 M899F | hNav1.7 M899I | hNav1.7 M899L | hNav1.7 M899V | GFP |
| --- | --- | --- | --- | --- | --- | --- | --- | --- |
| **Current densities (pA/pF)** | 490.3±96.4 (n=35) | 328.4±80.7 (n=26) | 250.4±74.0 (n=22) | 340.0±140.2 (n=20) | 26.1±6.4 (n=21) * | 182.6±36.6 (n=23) * | 114.6±19.4 (n=21) * | 18.0±1.9 (n=19) * |
| **$V_{1/2,act}$ (mV)** | -23.5±2.2 (n=35) | -25.5±2.0 (n=26) | -22.9±1.9 (n=22) | -20.6±2.2 (n=20) | - | -19.9±1.8 (n=23) | -26.6±2.3 (n=21) | - |
| **$Slope_{act}$** | 8.6±0.4 (n=35) | 8.3±0.3 (n=26) | 8.4±0.3 (n=22) | 8.3±0.7 (n=20) | - | 8.1±0.7 (n=23) | 9.3±0.7 (n=21) | - |
| **$V_{1/2,ssfi}$ (mV)** | -87.9±2.5 (n=17) | -87.5±2.2 (n=21) | -83.5±2.9 (n=16) | -88.0±2.5 (n=14) | - | -82.3±2.7 (n=17) * | -88.4±2.2 (n=18) | - |
| **$slope_{ssfi}$** | 6.6±0.8 (n=17) | 6.4±0.5 (n=21) | 7.7±0.8 (n=16) | 7.4±1.6 (n=14) | - | 7.1±1.0 (n=17) | 8.1±1.3 (n=18) | - |

Most gating parameters of activation and steady-state fast inactivation remained statistically insignificant between hNa_v_1.7 WT and the different substitutions (Figure 6C-F). The time to peak for hNa_v_1.7 M899C and M899L were observed in almost all voltage steps to be statistically significant and slightly slower than WT (p <0.05; Figure 6H), while the decay of fast inactivation was significantly slower for M899L in some depolarizing voltages when compared to WT (p <0.05; Figure 6I).

Overall, the experimental data largely supported the *in-silico* prediction that outer-pore bottlenecking is a major determinant of current density, whereas the various substitutions exerted minimal effects on other gating parameters.

## Discussion

Our study provides functional evidence for the pathogenicity of the CIP-linked hNa_v_1.7 M899I variant, causing a complete abolishment of Na_v_1.7 function in HEK293t cells without disrupting channel trafficking. We predict by coarse-grained MD that the variant causes a collapse of the outer pore by reducing the pore radius around the outer vestibule and the volume between the outer vestibule and the selectivity filter, with the branching of the amino acid side-chain at this location being more important than the size or functional group. We validated these predictions using in vitro patch-clamp experiments.

### A gating deficient mutant without trafficking defects

The lack of measurable sodium currents of hNa_v_1.7 M899I presents an added challenge in determining the reason for its loss-of-function. The lack of current can be either due to a lack of Na_v_1.7 M899I in the membrane due to trafficking defects or due to aberrant gating of hNa_v_1.7 M899I present in the membrane. In Na_v_1.1, many trafficking-deficient variants have been shown to be rescued by techniques such as co-expression with beta subunits (38) or lowering of the incubation temperature (39). Co-expressing Na_v_1.7 M899I with β2 or lowering the incubation temperature from 37°C to 30°C, however, did not significantly improve the current densities (Figure 2D).

The large size of Na_v_s renders them complicated for biochemical modifications and not many successful tagging attempts were reported so far. FLAG tags have been used to insert into various extracellular locations of hNa_v_1.5 to study membrane expression of LoF variants, such as in the extracellular loop between S1 and S2 in domain 1 (40) and the extracellular loop between S5 and S6 in domain 1 (41–43). However, the addition of the tag can alter the gating and activity of the tagged channels. For instance, the addition of various epitopes either extracellularly or intracellularly to hNa_v_1.5 caused drastic changes to the current densities and activation properties of the tagged channels (43). However, addition of the HA tag extracellularly to hNa_v_1.5 in one of the studies did not drastically alter surface expression (44). In our hands, insertion of an HA-tag did not alter expression and gating properties irrespective of the location (Figure S3). This allowed us to also show with a high degree of confidence that the hNa_v_1.7 M899I is expressed in the membrane (Figure 3B, white arrows).

The FLAG tag sequence “DYKDDDDK” contains many polar amino acids, while the HA-tag sequence “YPYDVPDYA” is relatively apolar. Extracellular loops in Na_v_s are typically charged and considered to influence the electrostatic field near the VSDs (45). Addition of a polar FLAG might have a drastic effect on the electrostatics than addition of more apolar tags such as HA. Thus, care must be taken about the composition of the tag to be added extracellularly. The lack of involvement in fast inactivation and having a long extracellular loop makes domain 1 an ideal region to add extracellular tags with minimal effect on channel gating. Our system provides a framework for efficiently and accurately showing membrane expression of LoF variants also in other Na_v_ subtypes and expression systems, including live cell staining of the cells to track real-time turnover of the channels.

### Lack of DN effects and the role of dimerization

Na_v_s, including Na_v_1.7, have been shown to dimerize and gate in a coupled manner. For example, the pain-causing variant Na_v_1.7 p.A1632E was shown to exert its GoF phenotype onto WT channels through dimerization when co-expressed in HEK293 cells (5). LoF variants in both Na_v_1.5 and Na_v_1.7 tend to exert DN effects, whereby a 50% co-expression of the non-function LoF variant with WT reduces the current densities by almost 75% (3, 5).

Our results on co-expression of hNa_v_1.7 M899I with hNa_v_1.7 WT showed a linear correlation to the amount of non-functional M899I expressed (Figure 2F-G), rather than the binomial relation typically observed for DN effects (3). This suggested a lack of DN effects in hNa_v_1.7 M899I. A Na_v_1.7 variant p.R896Q linked to CIP and located three amino acids downstream of M899I was shown to cause DN effects via dimerization (5). However, this effect was discussed as being caused by improper membrane trafficking of the LoF variant which would then dimerize with functional WT channels and prevent trafficking of the dimers. Although a recent study functionally characterized membrane-trafficking Na_v_1.7 LoF variants, they did not test for potential DN effects of these variants when co-expressed with WT channels (10).

Our results also highlighted the fully conserved nature of this residue, as the LoF was also observed when equivalent residues in Na_v_1.2 and Na_v_1.5 were mutated to either isoleucine or threonine (Figure 2E). This most likely suggests a conserved mechanism across the subtypes and allows us to draw comparisons to studies on DN effects of other Na_v_ subtypes. Many Na_v_1.5 LoF variants linked to Brugada syndrome have been characterized to cause DN effects, while a few variants do not (46). For example, the Na_v_1.5 p.R878C, which is a LoF variant of the Na_v_1.7 R896-equivalent residue, showed expression at the plasma membrane (47) and did not show DN effects (48). Another Na_v_1.5 LoF variant p.R893C, 11 amino acids upstream of the Na_v_1.7 M899-equivalent residue, also does not show DN effects and expresses in the membrane (49). The location, membrane expression and lack of DN effects of these few examples match well with what is observed with hNa_v_1.7 M899I. It could be that the lack of DN effects may be tied to either the variants being located in the PM of domain 2 or the variants having membrane expression similar to WT. However, a lack of DN effects may not necessarily mean a lack of dimerization. A recent study showed that the direct interaction of the α-subunits of Na_v_s may be independent from the coupling of their gating via secondary proteins like the 14-3-3 (50). Another study investigating a patient harboring compound heterozygous variants in Na_v_1.5 showed that the functional variant p.833R dimerized with WT channels while lacking dominant negative effects or coupled gating (51). The study argued this could be due to the disruption of the interface between the VSD and PM by p.G833R. This might also be likely for M899I channels – they may dimerize but lack the ability to gate in a coupled manner, most likely due to their location in PM2 potentially interfering with the interface between PM2 and VSD1.

All electrophysiological characterizations in this study were done in HEK293t cells. While they give us a good picture of the pathogenicity of the variant, it is not possible to make a direct causal link to the clinical phenotype. More physiological systems such as induced pluripotent stem cell-derived sensory neurons of the patient can help us in future studies to obtain a more complex picture of how these variants along with other factors can lead to CIP in the patient.

### Outer pore collapse as a possible patho-mechanism for loss of function

The outer pore has increasingly been shown to drive the process of ion conduction into the central cavity. Sodium ions are fully hydrated in bulk, usually coordinated by five to six water molecules. The SF is a very important region that imparts selectivity to Na_v_s for sodium ions and coordinates and partially dehydrates the sodium ion to enable passage towards the central cavity (52, 53). The residues in the OV have also been implicated in the coordination of the ion towards the SF (53, 54), also potentially acting as a beacon attracting these ions in a fully hydrated state towards the SF due to the highly electronegative nature of this region (55).

We observe a bottlenecking occurring between the SF and OV in hNa_v_1.7 M899I, as shown by the reduced volume in this pathway and pore radius close to the OV (Figure 4A-D). This bottlenecking reduces the diameter along this region and at the level of the OV to values much lower than the hydrated sodium ions. The radii seen for hNa_v_1.7 M899I at the outer vestibule are close to the ionic radius of a bare sodium ion (1.04A°). This would indicate that the fully hydrated sodium ion must undergo an energetically unfavorable dehydration of all its water molecules at the level of the OV even before it reaches the SF. This would make it unlikely for sodium to reach the SF and permeate further intracellular through the pore.

The collapsed outer pore may also signify the occupation of a slow inactivated state. A recent cryo-EM structure of Na_v_Rh showed a disruption of sodium ion conduction by dilation of the selectivity filter region (56). The literature, while obtained from non-mammalian channels, hints at the important role and sensitivity of outer pore geometry in deciding the meta-stable states that channels occupy. Cryo-EM structures of hNa_v_s, including the one used in the study, are still dominated by fast inactivated states due to methodological limitations. Obtaining cryo-EM structure of hNa_v_s in other gating states like the slow inactivated state can help us form better correlative insights to connect ion conduction to outer pore geometry.

The M899 residue is located close to the interface between VSD1 and PM2. This interface has been shown to be important in controlling the dynamics of VSD1, which is important for activation of the channel (57). It is possible that the residue could also disrupt this interface, resulting in inefficient dynamics of VSD1 and subsequent low activation probabilities. Further support for this hypothesis arises from previous observations that LoF variants may desynchronize transmembrane-helix rotations and ultimately decouple the VSD–PD interface (Xenakis & Lampert, 2025). This is, however, hard to assess with unbiased CGMD. Coarse-grained MD with Martini allows for longer, computationally inexpensive simulations relative to all-atom MD. This comes at the expense of resolution of the protein. Coarse-graining results in the loss of secondary and tertiary structure information. While elastic networks can circumvent this, it restricts the conformational space and therefore does not allow for observing large transitions such VSD movements or shifting from one state to another. Other computationally intensive techniques like umbrella sampling or metadynamics, allow for a larger conformational sample space and could give more insights into such mechanisms.

When using amino acid substitutions in position M899 we observed alteration of the extent of the outer pore bottleneck which seemed to be in line with the current densities observed for these variants (Figure 6). The linear shape of the amino acid might be crucial to maintaining packing of this region, with branching closest to the linear chains having a reduced effect on channel functioning. The M899, though fully conserved, seems to be robust to being replaced by various hydrophobic amino acids except for isoleucine. This region in the outer pore was shown to be a cluster for LoF variants to occur (58).

The robustness of M899 to substitutions also raise the question as to whether equivalent residues in other domains are equally robust to mutations. The substitution of M899-equivalent residues in Na_v_1.2 and Na_v_1.5 with threonine or isoleucine also resulted in a complete loss of function. It is thus highly likely that the mechanisms described in this study are applicable in a generalized manner across Na_v_ subtypes in various mammalian species. Further experiments in other Na_v_ subtypes could help us understand how the pore geometry of Na_v_s is maintained and where the sensitivity of the region arises from in a generalized manner. This in turn allows us to better predict the pathogenicity of Nav variants with unknown significance in the outer pore region tied to LoF-like clinical phenotypes in patients. Our study highlights a potential mechanism for LoF that helps understanding the basic mechanistics of Na_v_ gating, the mutational sensitivity of the channel structure and may help to reverse engineer efficient Na_v_ modifiers to reverse their hyper- or hypo-functionality.

## Conclusion

Our study characterizes a previously identified loss-of-function (LoF) variant Na_v_1.7 p.M899I linked to congenital insensitivity to pain. Using extracellular tagging approaches and whole-cell patch clamp, we show that the variant traffics to the plasma membrane yet conducts no detectable sodium currents. Coarse-grained simulations predict a bottlenecking of the outer pore, with graded effect for substitution of M899 with other hydrophobic residues. Patch clamp recordings follow the predicted rank order and correlate well with measured current densities. Based on our results, in addition to the conserved LoF phenotype across Na_v_ subtypes, we hypothesize a generalized mechanism for LoF variants in Na_v_s that collapse the outer pore and block ion conduction. Whether such a collapse could represent a unique gating state of the channel is an avenue to be further explored. Our work also highlights the ability of combined *in-silico* and *in-vitro* methods to reliably characterize Na_v_ variants and in turn better understand how these variants are linked to the disease phenotypes.

## Data Availability

The raw data and analysed datasets generated during the current study are available from the corresponding author on reasonable request.

## Author contributions

V.S.B.E and A.L designed the research and administered the project. V.S.B.E performed the whole-cell patch-clamp experiments, coarse-grained molecular dynamics simulations and structural visualization; wrote the code used to run the simulations and analyse the data; curated and analysed the data; and generated all figures. E.T.O. contributed to the whole-cell patch clamp experiments, data analysis and visualization of analysed data. A.N. and Y.L. performed immunostaining experiments shown in main text and supplementary material respectively. P.H. assisted with cell culture and generation of plasmids. A.B. and S.D.D. generated stable cell lines used for whole-cell patch clamp recordings. R.H. established the methodology for stable cell line generation. A.L and R.H provided the resources and funding for this work. A.L. and R.H. supervised with work with V.S.B.E playing a supporting role. V.S.B.E and A.L wrote the original draft. V.S.B.E, A.L., R.H., A.N., P.H., Y.L. and S.D.D. reviewed and edited the manuscript.

## Acknowledgements

We would like to acknowledge the help provided by Prof. Dr. Giulia Rossetti and Simone Albani for their advice about the CGMD workflows. We would also like to acknowledge the guidance of Dr. Markos Xenakis with utilizing entropy measurements such as KL-divergence and Dr. Jannis Körner for his input with respect to whole-cell patch clamp techniques. The authors would like to disclose that Claude AI was used in part to help in the creation of the custom analysis codes in Python. Microsoft Copilot was used for gentle language editing. This work was funded by DFG, German Research Foundation 363055819/GRK2415 (AL), DFG, German Research Foundation 368482240/GRK2416 (AL), DFG, German Research Foundation LA 2740/6-1 (AL). AL and RH were also supported by grants from the Interdisciplinary Center for Clinical Research within the Faculty of Medicine at the RWTH Aachen University IZKF (TN1-1/IA 532001 and TN1-5/IA 532005 respectively).

