## Supplementary Information for "Outer Pore Collapse as a Potential Mechanism of Partial Loss of Pain in Nav1.7 M899I"

### Supplmentary Information for:

**This PDF includes:**

**Figure S1 – Immunostaining of hNav1.7 M899I**

**Figure S2 - Electrophysiological characterization of extracellularly HA-tagged hNav1.7**

**Figure S3 - Root mean square deviations of simulated hNav1.7 WT and variants**

**Figure S4 - Root mean square fluctuations of simulated hNav1.7 WT and variants**

**Table S1 - Current densities and gating parameters of HEK293 Flp-in™ cells transiently transfected with either hNav1.7 WT or various substitutions of the M899 residue (M899I, M899A, M899C, M899F, M899L or M899V).**

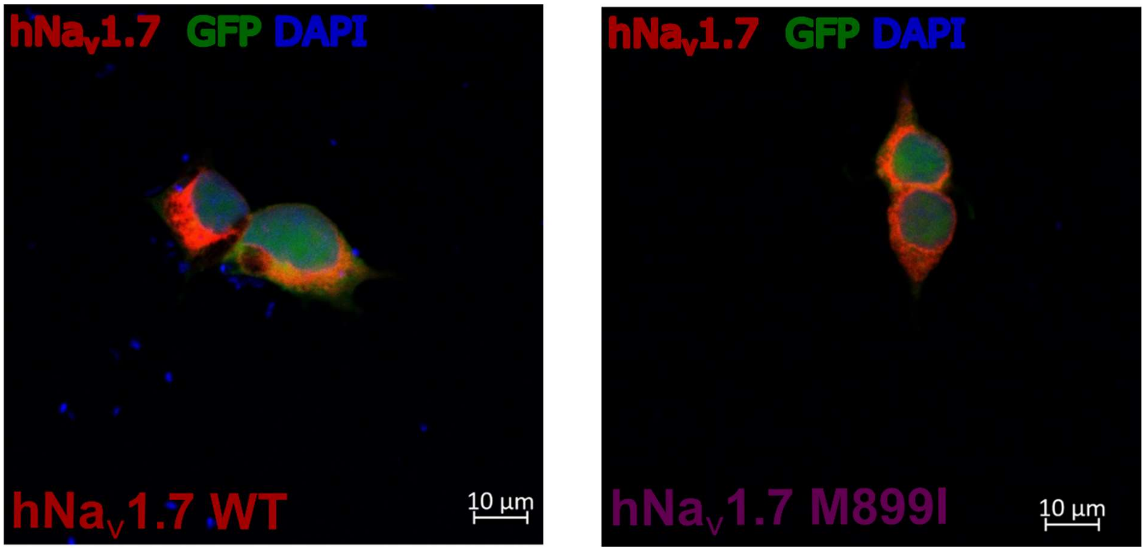


**Figure S1: Immunostaining of hNa$_v$1.7 M899I.** Confocal imaging of HEK293t cells transfected with either hNa$_v$1.7 WT (left) or hNa$_v$1.7 M899I and immunostained with antibodies targeting hNa$_v$1.7. The reporter protein GFP is observed using a green signal, while blue signal represents the DAPI staining of the nucleus. Scale bar is shown in the bottom right corner. Due to the reduced specificity of the hNa$_v$1.7 antibody, endogenous expression of Na$_v$1.7 in HEK293t cells and the need for membrane permeabilization of the transfected cells, clear discernment cannot be made between intracellular channels and channels in the membrane bilayer.

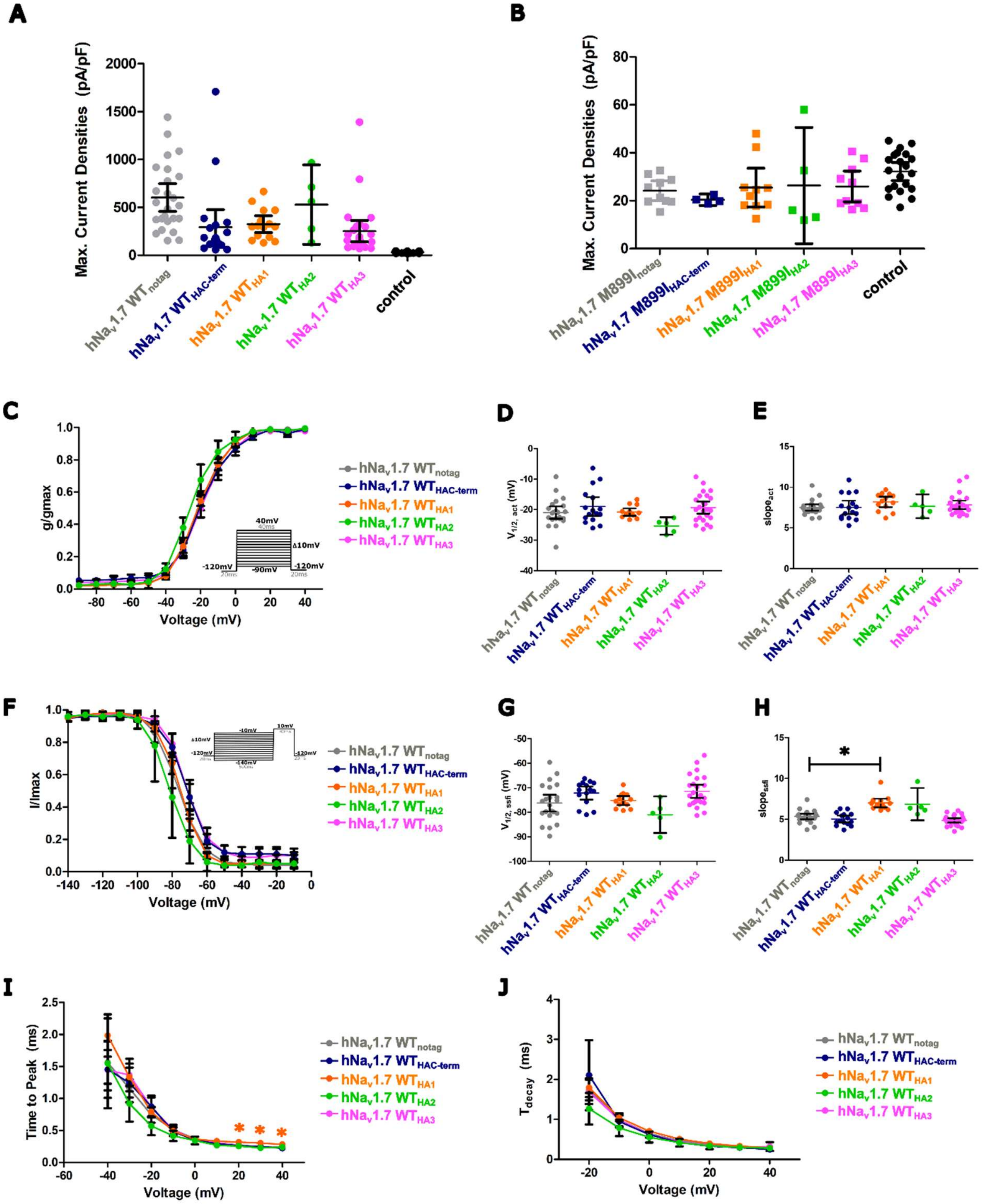


**Figure S2: Electrophysiological characterization of extracellularly HA-tagged hNa$_v$1.7. A)** Dot plot of the maximal current densities of HEK293t cells stable expressing hNa$_v$1.7 WT without (notag) or with HA tags inserted intracellularly (HAC-term) or extracellularly (HA1, HA2 and HA3). Addition of the HA tag do not alter the current densities of hNa$_v$1.7 WT. **B)** Dot plot of the maximal current densities of HEK293t cells stable expressing hNa$_v$1.7 M899I without (notag) or with HA tags inserted intracellularly (HAC-term) or extracellularly (HA1, HA2 and HA3). Irrespective of the presence or location of the HA-tag, hNa$_v$1.7 M899I activity is indistinguishable from the control. **C)** Normalized conductance curves measured using the protocol in inset, **D)** $V_{1/2,act}$ and **E)** slope$_{act}$ for hNa$_v$1.7 WT without (notag) or with HA tags inserted intracellularly (HAC-term) or extracellularly (HA1, HA2 and HA3). All activation parameters remained similar, irrespective of the presence and location of the HA-tag. **F)** Normalized current curves measured using the protocol in inset, **G)** $V_{1/2,ssfi}$ and **H)** slope$_{ssfi}$ for hNa$_v$1.7 WT without (notag) or with HA tags inserted intracellularly (HAC-term) or extracellularly (HA1, HA2 and HA3). Most steady-state fast inactivation parameters remained similar, irrespective of the presence and location of the HA-tag. **(I)** Time to peak and **(J)** onset of fast inactivation kinetics for hNa$_V$1.7 WT without (notag) or with HA tags inserted intracellularly (HAC-term) or extracellularly (HA1, HA2 and HA3). Very minor changes in the kinetic properties were observed. * $p<0.05$. The control are pooled data from HEK293t cells stably expressing hNa$_V$1.7 plasmids without the addition of doxycycline. The error bars represent the 95% confidence interval of the mean.

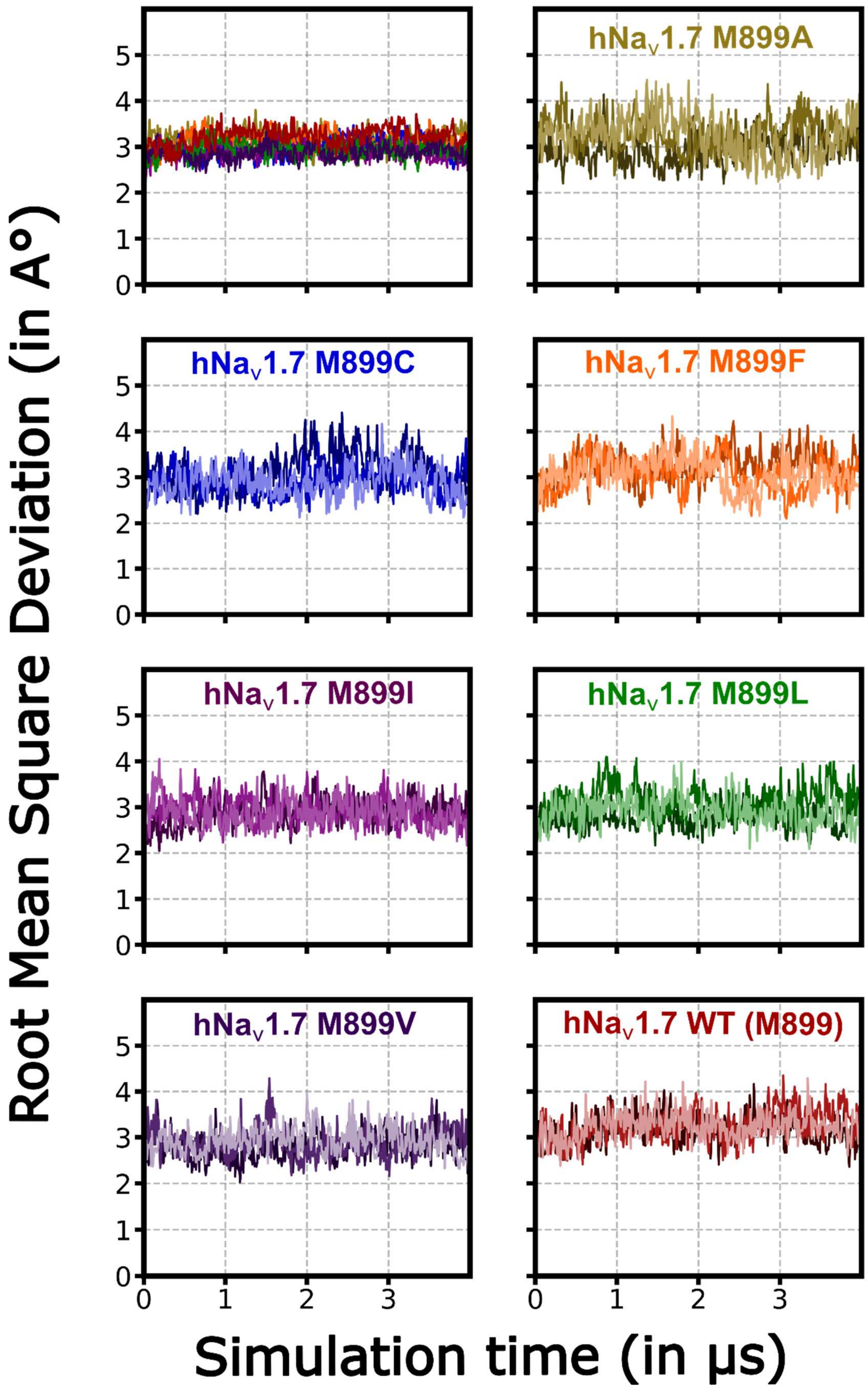


**Figure S3: Root mean square deviations of simulated hNa$_V$1.7 WT and variants.** Root mean square deviations (RMSD) of the WT or variants obtained from post-processed simulation frames using gmx rms. RMSD is plotted over the simulation time.

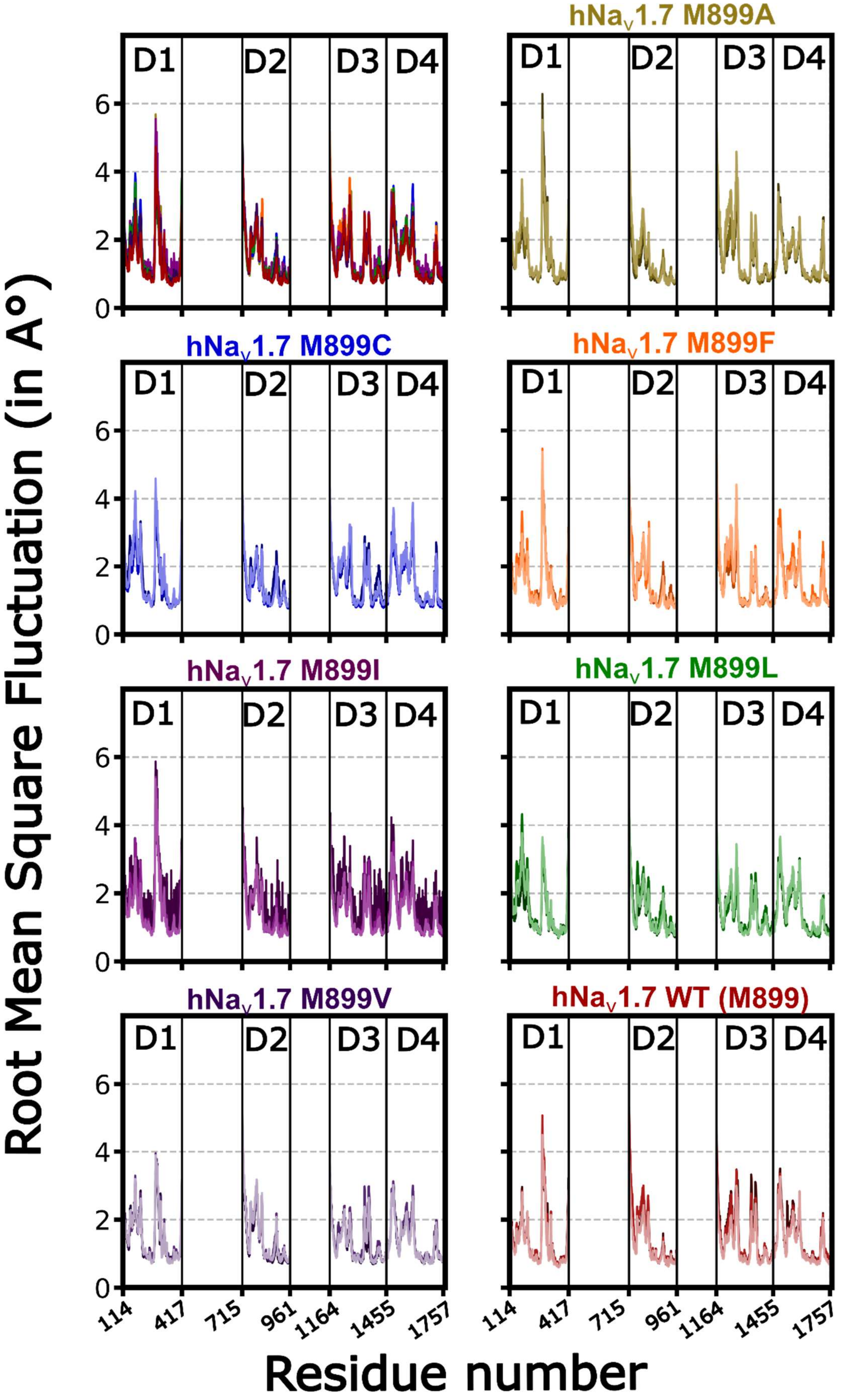


**Figure S4: Root mean square fluctuations of simulated hNa$_V$1.7 WT and variants.** Root mean square fluctuations (RMSF) of the WT or variants obtained from post-processed simulation frames using gmx rmsf. RMSF is plotted over the simulation time.

**Table S1 – Current densities and gating parameters of HEK293 Flp-in™ cells transiently transfected with either h$Na_v$1.7 WT or various substitutions of the M899 residue (M899I, M899A, M899C, M899F, M899L or M899V).** GFP transfection acts as negative control. Gating parameters include the $V_{1/2,act}$, $slope_{act}$, $V_{1/2,ssfi}$ and $slope_{ssfi}$. Data are always presented as mean±95% confidence interval (95% CI) of the mean. n represents the number of cells/data points. *p <0.05 when compared to h$Na_v$1.7 WT.

| | **h$Na_v$1.7 $WT_{notag}$** | **h$Na_v$1.7 $WT_{C\text{-}term}$** | **h$Na_v$1.7 $WT_{HA1}$** | **h$Na_v$1.7 $WT_{HA2}$** | **h$Na_v$1.7 $WT_{HA3}$** | **Control** |
|---|---|---|---|---|---|---|
| **Current densities (pA/pF)** | 602.4±144.8 (n=25) | 294.7±182.2 (n=15) | 325.3±87.8 (n=15) | 529.7±415.2 (n=5) | 252.5±112.2 (n=26) | 32.2±3.7 (n=21) * |
| **$V_{1/2,act}$ (mV)** | -21.0±2.0 (n=22) | -19.1±2.0 (n=16) | -20.9±1.2 (n=13) | -25.4±2.9 (n=5) | -19.4±2.0 (n=24) | - |
| **$Slope_{act}$** | 7.5±0.4 (n=22) | 7.5±0.8 (n=16) | 8.2±0.6 (n=13) | 7.6±1.5 (n=5) | 7.8±0.5 (n=24) | - |
| **$V_{1/2,ssfi}$ (mV)** | -76.2±3.4 (n=22) | -72.2±2.7 (n=15) | -75.2±1.8 (n=13) | -81.0±7.5 (n=5) | -71.4±2.7 (n=24) | - |
| **$slope_{ssfi}$** | 5.4±0.3 (n=22) | 5.0±0.4 (n=15) | 7.0±0.5 (n=13) * | 6.9±2.0 (n=5) | 4.9±0.3 (n=24) | - |
